# Finger stability in precision grips

**DOI:** 10.1101/2021.12.04.470276

**Authors:** Neelima Sharma, Madhusudhan Venkadesan

## Abstract

Stable precision grips using the fingertips are a cornerstone of human hand dexterity. Occasionally, however, our fingers become unstable and snap into a hyper-extended posture. This is because multi-link mechanisms, like our fingers, can buckle under tip forces. Suppressing this instability is crucial for hand dexterity, but how the neuromuscular system does so is unknown. Here we show that finger stability is due to the stiffness from muscle contraction and likely not feedback control. We recorded maximal force application with the index finger and found that most buckling events lasted less than 50ms, too fast for sensorimotor feedback to act. However, a biomechanical model of the finger predicted that muscle-induced stiffness is also insufficient for stability at maximal force unless we add springs to stiffen the joints. We tested this prediction in 39 volunteers. Upon adding stiffness, maximal force increased by 34±3%, and muscle electromyography readings were 21±3% higher for the finger flexors (mean±standard error). Hence, people refrain from applying truly maximal force unless an external stabilizing stiffness allows their muscles to apply higher force without losing stability. Muscle recordings and mathematical modeling show that the splint offloads the demand for muscle co-contraction and this reduced co-contraction with the splint underlies the increase in force. But more stiffness is not always better. Stiff fingers would interfere the ability to passively adapt to complex object geometries and precisely regulate force. Thus, our results show how hand function arises from neurally tuned muscle stiffness that balances finger stability with compliance.

## I. INTRODUCTION

Precision grip, as the name implies, is the precise and stable application of fingertip forces. In this grip style, the fingers are relatively stationary while the fingertips exert force (Napier, 1956). A stable precision grip played a key role in the evolution of human hand dexterity (Karakostis *et al*., 2018; Kivell, 2015; Marzke, 1997, 2013). But the inherent mechanics of multi-link chains make the fingers prone to many types of instabilities when the fingertip experiences forces (Hogan and Buerger, 2018; Murray *et al*., 2017). The nervous system often masks these instabilities by using a lifetime of learned control strategies. So we rarely witness them in everyday experience. Understanding how the nervous system suppresses these instabilities is needed to explain hand function and its loss due to disease or aging.

Instabilities that arise when pushing on surfaces can be categorized as those affecting the tip where the force is applied (Bicchi and Kumar, 2000; Hogan and Buerger, 2018; Murray *et al*., 2017; Okamura *et al*., 2000; Rancourt and Hogan, 2001; Whitney, 1987), or the internal degrees of freedom associated with posture (Bunderson *et al*., 2008; De Groote *et al*., 2017; Klimchik *et al*., 2015). Tip instabilities are particularly severe when a stiff finger or limb makes contact with a rigid surface (Hogan and Buerger, 2018). When feedback control is used to precisely apply tip forces, the fingertip’s position in space may become unstable and start to oscillate, which also destabilizes the applied force (Hogan and Buerger, 2018; Whitney, 1987). One strategy is to increase the compliance of the finger or limb (Akella and Cutkosky, 1989; Hanafusa and Asada, 1977; Kao *et al*., 1997; Mason, 1981). Such stiffness control and its generalization to impedance control in dynamic contexts (Hogan and Buerger, 2018) have proven quite effective in controlling contacts in robots.

Postural stability of the internal degrees of freedom has received considerably lesser attention than tip stability, and has only been studied in models (Bunderson *et al*., 2008; De Groote *et al*., 2017) or robots (Klimchik *et al*., 2015). Kinematic chains with many internal degrees of freedom are prone to buckle and lose postural stability under external compressive forces (supplement §S1.1 and §S1.2), analogous to the buckling of slender columns. Consider pushing a rigid surface with the index fingertip (figure 1), and focus on the mechanics within the plane of the finger. In this setting, the index finger has three internal degrees of freedom between the knuckle and the tip. When the tip does not slip, it is subject to two translational constraints within the finger’s plane. Thus the finger is kinematically underdetermined by one degree. It is this degree of freedom that could become unstable.

**FIG. 1.**
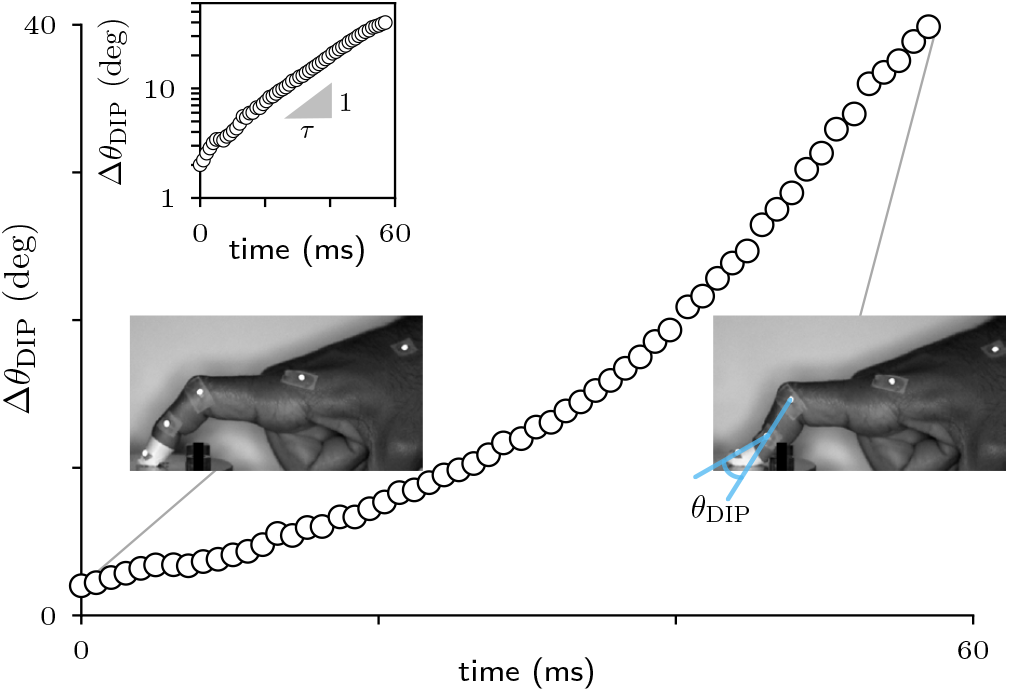
Buckling of the index finger joints. Sample trial showing the change in the angle of the distal interphalangeal joint (DIP), Δ*θ*_DIP_. Every fifth sample is plotted for clarity (black circles). **inset:** Linear-log plot of the exponential growth in DIP angle. The time-constant *τ* for the unstable growth in Δ*θ*_DIP_ is found using the slope. For this trial, *τ* = 20 ms.

To suppress postural instabilities, humans appear to use a conflicting strategy to that of stable force control, namely to make the limb stiffer. Potential instabilities of limb or finger posture when applying contact force have not been studied much, but a related behavior of stabilizing the posture of a handheld tool has been investigated previously (Rancourt and Hogan, 2001). When using a tool like a handheld drill, more force applied on the wall makes the orientation of the drill more susceptible to becoming unstable (Rancourt and Hogan, 2001). Rancourt and Hogan (2001) found that hand stiffness is critical for stabilizing the drill. The nervous system uses stiffening as a strategy for postural stability in other contexts as well, such as dealing with unstable environmental dynamics when moving the arm (Burdet *et al*., 2001) or the destabilizing effects of motor noise (Selen *et al*., 2009). Higher stiffness, which is harmful for tip stability under force feedback control, may be what stabilizes the internal degrees of freedom of our fingers and limbs. But the role of stiffness remains debated and unresolved in several contexts involving postural stability. Examples include standing in humans (De Groote *et al*., 2017; Peterka, 2002) and cats (Bunderson *et al*., 2008), and arm (Selen *et al*., 2009) and finger movements (Venkadesan and Valero-Cuevas, 2008). We presently lack studies to tease apart the role of stiffness versus other strategies such as feedback control for maintaining postural stability during contact.

In this paper, we investigate postural stability of our fingers during maximal force application by using the index finger as a representative example. We study flexed postures where musculotendon tissues support the joint loads because of its implications for understanding forceful precision grasps (Karakostis *et al*., 2018; Kivell, 2015; Marzke, 2013; Marzke and Marzke, 2000), and do not consider hyper-extended postures due to potential of injury when producing large tip forces (Marco *et al*., 1998; Schweizer, 2001; Vigouroux *et al*., 2006). Our approach is inspired by the study of Rancourt and Hogan (2001), who used maximal force tasks to probe the neural strategy that stabilizes the posture of a handheld drill. The central idea is to challenge the nervous system by making the internal mechanical response as unstable as possible. Because buckling-type instabilities are generally more severe at higher forces, we examined how the fingertip’s maximal force is affected by external modifications to the finger that alter stability. Moreover, the muscle activation pattern that people use at sub-maximal fingertip force is a linearly scaled version of the pattern that they use at maximal force (Valero-Cuevas, 2000; Valero-Cuevas *et al*., 2009; Venkadesan and Valero-Cuevas, 2008). So, understanding stability at maximal force may also help us understand the properties at sub-maximal forces. In a series of experimental and mathematical studies, we show that feedback control alone cannot stabilize the postural instability during maximal tip force application, and people rely on the spring-like properties of muscles to suppress the instability.

## II. RESULTS

### A. Postural instability of the index finger

We first conducted a study with nine volunteers to assess the severity of the buckling instability in the index finger (see Methods). They were instructed to apply the largest normal force possible on a rigid surface, with no explicit instruction about stability. We asked them to repeatedly try to push harder until we recorded 33 instances of postural instabilities. The instability manifested as a sudden change in the finger’s posture where one of the three finger joints ended up in a hyper-extended angle. The distal interphalangeal joint (DIP) buckled most frequently, in 28 out of 33 instances that we captured (figure 1, movie 1). So, we used the DIP angle to analyze the temporal characteristics of the buckling event. The DIP joint angle grew exponentially in the trials (R^2^ > 0.9). The time-constant was smaller than 45 ms in 19 out of 28 trials, and never exceeded 80 ms (figure 1, table S3).

Neural feedback control alone cannot stabilize such rapid instabilities because the nerve conduction latency for the round-trip from the hand to the spinal cord exceeds 45 ms, and the fastest sensory-driven finger response is usually timed at 65 ms or more (Johansson and Birznieks, 2004). Despite these latencies, the finger was stable for most of the trials (40 out of 63 trials were stable in 7 subjects, the number of non-buckling events were not recorded in the remaining two). Thus, we postulate that muscle contraction and the joint stiffness it induces is responsible for stability. Muscles are intrinsically stiffer when producing more force, a property known as short-range stiffness (Cui *et al*., 2008; Rack and Westbury, 1974). Therefore, the harder someone pushes with the fingertip, the higher the muscle and overall finger stiffness ((Hajian and Howe, 1997), supplement §S1.1). So, stability could just be a byproduct of the muscles contracting to produce force. Alternatively, the need to remain stable may constrain the maximum exertion of fingertip force. We performed additional analyses and experiments to find out whether stability is a byproduct or a constraint.

### B. Stability at maximal force

We investigated postural stability using a previously developed detailed anatomical model of the index finger (Valero-Cuevas *et al*., 1998; Venkadesan and Valero-Cuevas, 2009, supplement §S1.1). The finger is modeled as a three-link, planar kinematic chain, and driven by seven muscles (figure 2a). The tip is constrained to not translate to capture the absence of fingertip slip, but can freely rotate (Venkadesan and Valero-Cuevas, 2009). Muscle activations are specified by a normalized 7D vector 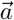 with values between 0 and 1, which govern both muscle force and stiffness (Bunderson *et al*., 2008; Cui *et al*., 2008, supplement §S1.1). Activating the muscles drives the finger’s joints with torques 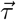, applies a fingertip force 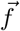, and induces stiffness K at the joints.

**FIG. 2.**
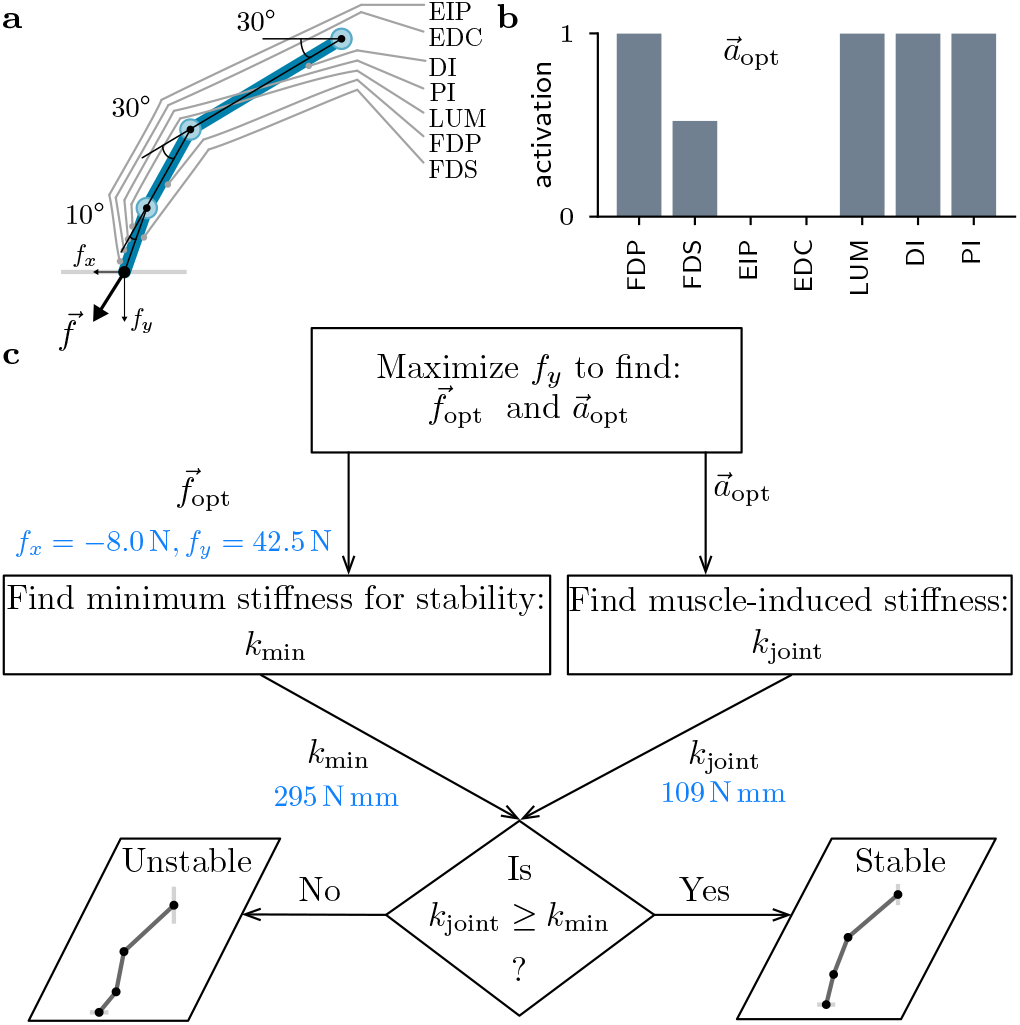
Modeling study to test if stability is a byproduct of fingertip force. **a,** Schematic of a planar model of the index finger that maintains contact at the fingertip and is driven by seven muscles. **b,** The optimal activation pattern 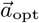 that maximizes the vertical component of the fingertip force at a fixed posture 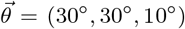. This is the posture used in subsequent experiments in this paper. **c,** The decision tree to test whether muscle-induced stiffness leads to stability when the activation pattern is chosen solely to maximize fingertip force. The computed force and stiffnesses at 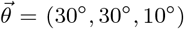 are in blue. The finger is unstable at the maximal force because *k*_joint_ < *k*_min_.

The planar finger model has only one unconstrained degree of freedom. So the constrained dynamics of the finger are defined by a projection of the finger’s dynamics onto the null-space of the constraints that are imposed on the fingertip (supplement §S1.1). The orthonormal basis vectors of the null-space are expressed as columns of the null-space matrix P, which in the case of the single degree of freedom finger reduces to a single null-space vector. Thus, the 3 × 3 stiffness matrix K associated with the multi-link finger reduces to a scalar stiffness *k*_joint_ = P^T^KP when projected onto the finger’s unconstrained degree of freedom. In the absence of feedback control, stability requires the muscle-induced joint stiffness *k*_joint_ to exceed a minimum threshold *k*_min_ that depends on the tip force (supplement §S1.2),

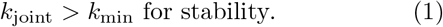

We computed the optimal muscle activation pattern 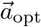 that maximizes the vertical force without any constraints imposed on stability (figure 2b, supplement §S1.3). This activation produces a tip force 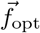 and joint stiffness *k*_joint_ because of muscle’s short-range stiffness. This stiffness *k*_joint_ was compared with the minimum stiffness *k*_min_ that is needed for stability at the maximal force 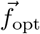 (figure 2c, supplement §S1.2). For the same posture as the experiments (30°, 30°, 10°), we found *k*_joint_ = 109 N mm and *k*_min_ = 295 N mm. So, the finger is unstable for the activation pattern that maximizes tip force (figure 2). We also examined all possible postures that do not hyper-extend the joints while maintaining the same tip position as (30°, 30°, 10°). None of those postures were stable at the maximal force (figure S1). Therefore, stability does not automatically arise as a byproduct of force application.

Once the finger becomes unstable, the posture grows along the unstable mode specified by the null-space vector P. For the posture (30°, 30°, 10°), the null-space vector is P = (0.06-0.49 0.87)^T^. The third, DIP component of the vector is the biggest in magnitude, indicating that the largest change in joint angles would occur at the DIP joint. This behavior is consistent with the recorded buckling events in human subjects (section II.A) and numerical simulation of the nonlinear equations of the finger model (supplement §S1.4, movie 2).

The results support the hypothesis that stability rather than muscular capacity constrains the maximal voluntary fingertip force. Therefore, we predict that people should be able to produce more force if their finger is externally stiffened to reduce or eliminate the postural instability. We tested this prediction in experiments with volunteers.

### C. Effect of externally stiffening the finger

We instructed 39 consenting volunteers to stably apply the largest force they could with their index fingertip against a rigid surface (see Methods). To test the prediction of the model, we compared the maximal force of a free index finger with trials where we externally stiffened the finger by attaching a custom-molded thermoplastic splint. Motivated by the large DIP component of the unstable mode, we used two splint designs (figure 3a). One that stiffened the DIP and the PIP joints (2J splint), and another that only stiffened the PIP joint (1J splint). Because the finger has only one net degree of freedom, both splint designs would stiffen the finger, with lesser stiffness induced by the 1J splint. We recorded, smoothed, and processed the fingertip force and surface electromyograms (EMG) of the *flexor digitorum profundus* (FDP) and *flexor digitorum superficialis* (FDS) from all 39 subjects and additionally recorded EMG of the *extensor digitorum communis* (EDC) in a subset of 16 subjects. The maximal force F_max_ is the maximum of the force trace that is smoothed using a 1-second window. The normalized maximal force *f*_max_ is the maximal force normalized by the maximal force from all the trials of a given subject. The flexor EMG measure EMG_flexors_ is a PCSA-weighted average of the RMS of the filtered and MVC-normalized EMG from the two flexors during the 1-second window where force is maximized (see Methods, figure 3b). Data from one subject was excluded because their finger repeatedly buckled without the splint and did not yield reliable measurements.

**FIG. 3.**
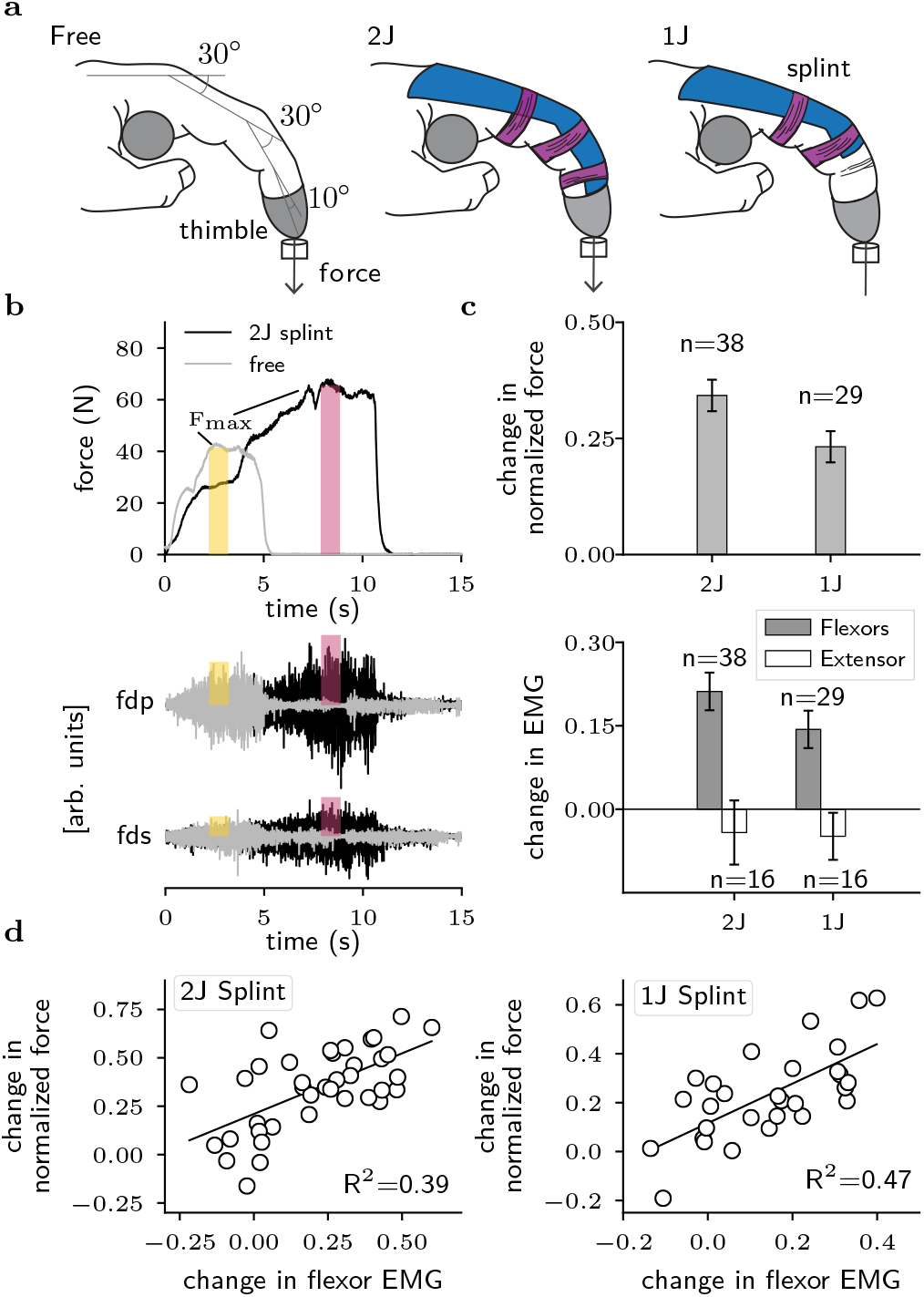
Maximal force upon stiffening the finger. **a,** Three conditions were tested at the posture (30°, 30°, 10°): no splint (free), 2-joint split (2J), or 1-joint splint (1J). **b,** For a sample subject, the shaded rectangles show the time-window when the maximal force occurred, pink for 2J and yellow for free, overlaid on the vertical force, and raw EMG recordings from FDP and FDS. EMG rectangles are scaled 6× for clarity, but the force rectangles are to scale. **c,** Change in the maximal normalized force *f*_max_, flexor EMG, and extensor EMG for the 2J and 1J conditions away from the free finger. The bars and whiskers show the mean and standard error, respectively. **d,** Scatter plots and regression fits of the change in EMG versus change in force between the splint and the free conditions, for the 2J and 1J conditions.

Presence of a splint significantly affected the normalized maximal force *f*_max_ (*F*_2,263_ = 144.57, *p* < 0.0001), and EMG_flexors_ (*F*_2,263_ = 40.22, *p* < 0.0001). The order of trials, free or splinted first, did not have a significant effect on *f*_max_ or EMG_flexors_ (*F*_1,218_ = 2.13, *p* = 0.15 and *F*_1,190_ = 0.46, *p* = 0.49, respectively). There was also no significant interaction between splint condition and the order of presentation of the trials on either *f*_max_ or EMG_flexors_ (*F*_2,68_ = 2.28, *p* = 0.11 and *F*_2,69_ = 2.36, *p* = 0.10, respectively). Relative to the free finger, the normalized maximal force significantly increased for the 2-joint and 1-joint conditions by Δ*f*_max_ = 0.34±0.03 and 0.23±0.03, respectively (mean±standard error, figure 3c, *p* < 0.0001 in both conditions). There was subject-to-subject variability in the magnitude of increase, but the force increased for all but three subjects with the 2-joint splint, and for all but one with the 1-joint splint. EMG from flexors also significantly increased for the 2-joint (*p* = 0.04) and 1-joint conditions (*p* = 0.04), by ΔEMG_flexors_ =0.21±0.03 and 0.14±0.03, respectively (mean±standard error, figure 3c). Statistically significant differences were not found between the two splint types either for normalized force or EMG from flexors.

The normalized force increase Δ*f*_max_ and ΔEMG_flexors_ are significantly correlated for both the 2-joint (R^2^=0.39, *p* < 0.0001) and 1-joint conditions (R^2^=0.47, *p* < 0.0001), despite the generally noisy nature of surface EMG measurements, showing that the increased tip force was because of higher muscle force (figure 3d). Detailed statistics and verification of assumptions are in supplement §S2.

We conclude that the nervous system refrains from producing truly maximal force. Upon stiffening the finger, especially the DIP joint in the 2-joint splint, the maximal force increased. This is consistent with the prediction that once stability is no longer a concern, higher force can be applied. The increase in flexor EMG activity with force indicates that the nervous system could tap into additional muscle force capacity, but only when the finger was externally stabilized. We next perform an analysis of co-contraction in order to test this idea on additional muscle force capacity with the splint and to understand the origin of inter-subject variability.

### D. Co-contraction and maximal force

The idea behind more force with the splint is that the splint provides stiffness for stability, thus allowing muscle co-contraction to decrease, which in turn allows for more force from the flexors that contribute substantially to tip force. To analyze changes in co-contraction with the splint, we recorded surface EMG from the extrinsic extensor EDC in a subset of 16 subjects, in addition to the extrinsic flexors FDP and FDS. The force and the EMGs are normalized per subject by the respective largest recorded value, and the aggregate EMG_flexors_ is defined as in §II.C. The extensor and flexors are approximate antagonists, implying that lesser co-contraction would manifest as a reduction in the ratio EMG_EDC_/EMG_flexors_ upon adding the splint. A one-way ANOVA with finger condition (free, 1J, or 2J) as the factor and the ratio EMG_EDC_/EMG_flexors_ as the dependent variable was significant (*F*_2,45_ = 3.80, *p* = 0.03). Post hoc contrasts show that the ratio is significantly smaller for the 2J splint (*p* = 0.049, Δratio = −0.43), and borderline for the 1J splint (*p* = 0.06, Δratio = −0.42) compared to the free finger (figure 4a, supplement table S10).

**FIG. 4.**
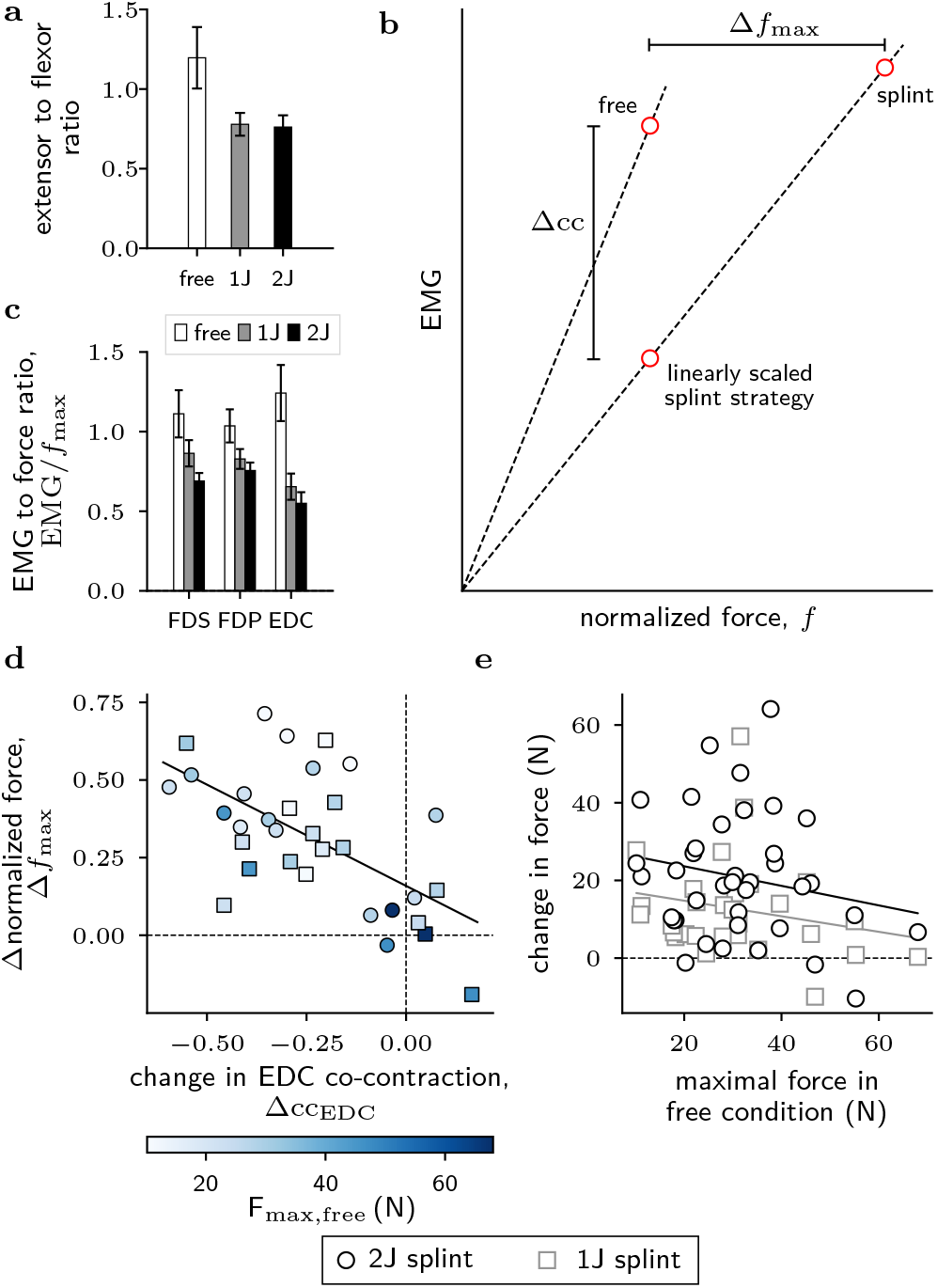
Co-contraction and maximal force. **a,** Ratio of extensor to flexor activity for the free, 1J, and 2J splint conditions (n=16). The bars and whiskers show the mean and standard error, respectively. **b,** Pictorial demonstration of the hypothesis that the free finger is co-contracted relative to the splinted condition, as seen by a steeper slope for the free finger compared to the splint condition in the normalized EMG-force space. **c,** EMG to normalized force ratios for the two flexors, FDS and FDP, and the extensor, EDC for the free, 1J and 2J splint conditions (n=16). The bars and whiskers show the mean and standard error, respectively. **d,** Scatter plot and regression fit of the change in EDC co-contraction versus the change in normalized force. The scatter plot is colored by the magnitude of the free finger’s baseline force (n=16). **e,** Scatter plot and regression fits of the free finger’s baseline force versus the change in force between the splinted and the free conditions for the 2J (black, n=38) and the 1J splints (gray, n=29).

However, the finger’s muscles are not organized as a simple uniarticular agonist-antagonist system. Therefore, producing joint torques to apply fingertip force in a specific direction leads to co-activation of even seemingly antagonistic muscles (Valero-Cuevas *et al*., 1998), and produces joint stiffness as a byproduct (§II.B, figure 2b). To account for these complexities and refine the analysis of co-contraction, we use the decomposition of the total EMG recorded for a muscle as one portion that contributes to fingertip force with stiffness as a byproduct, and another that lies in the null-space of the mapping from muscle contraction to fingertip force and thus contributes solely to finger joint stiffness but not the fingertip force. We term the latter component as co-contraction. The hypothesized mechanism for force increase with the splint is that the null-space component needed for stability is lowered upon adding the splint. Therefore, if the splint reduced the co-contraction component, the same force could be produced with lesser EMG, i.e. the ratio of EMG to force would be lesser with the splint (schematic of hypothesis in figure 4b). The one-way ANOVAs with finger condition as the factor and the EMG to force ratio as the dependent variable were significant for all three muscles (FDS: *F*_2,45_ = 4.29, *p* = 0.02, FDP: *F*_2,45_ = 3.69, *p* = 0.03, EDC: *F*_2,45_ = 9.79, *p* = 0.0003, figure 4c). Post hoc contrasts show that the 2J splint always led to a significant decrease in the EMG to force ratio, but the 1J splint led to a significant ratio decrease only for the EDC (supplement table S11). As seen from the optimization for tip force (§II.B, figure 2b), EDC has a nil or weak projection onto the tip force for the posture used in our study. So EDC activity is probably most strongly associated with finger stiffening. This understanding of EDC function is consistent with the results that its EMG was most sensitive to adding a splint, 1J or 2J, and responded by nearly halving in magnitude.

The co-contraction analysis could also help understand the inter-individual differences in force increase with the splint. It is possible that not all subjects increased their force output by the same relative magnitude when the splint was added because of differences in the extent of reduction in co-contraction. More important is the new understanding that the splint is simply a means to allow the subject to reduce their co-contraction and the force gain is hypothesized to be a result of the reduced co-contraction. To assess this, we extended the analysis of EMG to force ratio to assess how well the decrease in EDC contraction could explain the change in maximal voluntary force. The EDC is chosen as the primary muscle for the analysis because of its more direct role in finger stiffness and minimal or nil contribution to fingertip force. Informed by previous studies that showed that people linearly scale their EMG patterns as they vary their force (Valero-Cuevas, 2000; Valero-Cuevas *et al*., 2009; Venkadesan and Valero-Cuevas, 2008), we compared the free finger’s EDC EMG with a linearly scaled version of the splinted condition EDC EMG to estimate the excess EMG in the free finger relative to the splinted condition if the force were the same (figure 4b). The excess EDC co-contraction Δcc_EDC_ is defined as the difference in EMG between the linearly scaled splinted trial and the free trial according to,

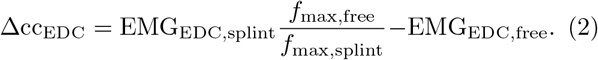

Lesser co-contraction would imply a negative Δcc_EDC_. A linear regression, with the change in normalized force Δ*f*_max_ as the dependent variable and Δcc_EDC_ as the regressor, showed significant correlation between co-contraction change and force change (*F*_1,30_ = 16.93, *p* = 0.0003, *R*^2^ = 0.36, slope±standard error= −0.65 ± 0.16, intercept±standard error= 0.15 ± 0.05, figure figure 4d). We also tested an alternative hypothesis that the force gain with the splint is minimal or zero in some subjects because they are not limited by stability to begin with and would exhibit high force capabilities even without the splint. We tested this using a regression with the non-normalized increase in force as the dependent variable and non-normalized force under free conditions as the regressor. This regression was not significant for either the 2J or 1J splints (2J: *F*_1,36_ = 1.48, *p* = 0.23, 1J: *F*_1,27_ = 1.34, *p* = 0.26, figure 4e). Furthermore, the change in co-contraction Δcc_EDC_ was not significantly correlated with the free finger’s maximal force (*F*_1,30_ = 3.31, *p* = 0.08), thus making it unlikely that differences in co-contraction reduction are simply because some subjects were strong to begin with.

The inter-individual variability in reduction in co-contraction may reflect differences in willingness to explore new areas of muscle contraction space that is unlike the normal experience. The baseline force was unable to explain the force difference showing that people who are strong to begin with could still produce more force with the splint, so long as they decreased their co-contraction. Surface EMG, that too from a partial subset of the muscles of the finger, is a potentially noisy measure of co-contraction but, as we find, with predictive power. The choice of the EDC that is informed by the modeling analysis probably helped detect the role of co-contraction in force production despite potential limitations of surface EMGs. Future studies could refine our results and probe learning and neuromotor exploration of the feasible space of strategies by using intramuscular readings from all seven muscles.

### E. Sub-maximal forces

The trade-off between co-contraction and maximal force suggest that at sub-maximal forces, the muscles may have the leeway to co-contract and modulate stiffness without affecting force. To investigate this, we used the finger model and numerically found activation patterns with minimum and maximum joint stiffness *k*_joint_ when applying a sub-maximal force (‘1’ and ‘4’ in figure 5a, supplement §S1.5, posture: 30°, 30°, 10°). Both these patterns produced the same horizontal force as the maximal solution (figure 2b) but just 9 N of force vertically, which is 4.68× lower than maximal. Because the force is sub-maximal, there is a four-dimensional nullspace for the mapping from muscle activations to fingertip force. All the null-space activation patterns apply the same joint torques and tip forces, but their stiffness would vary between those of ‘1’ and ‘4’.

**FIG. 5.**
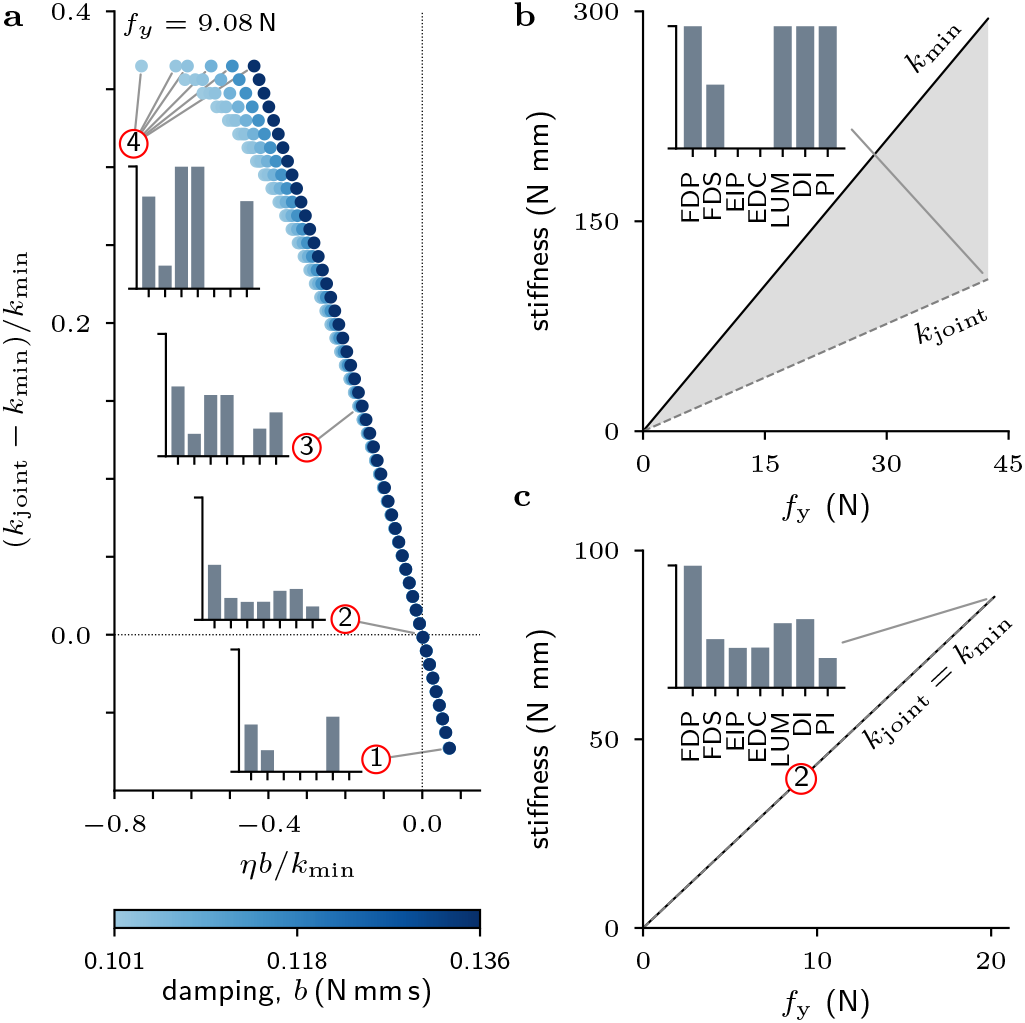
Muscle co-contraction, stiffness, and stability at sub-maximal force. **a,** Monte Carlo simulations densely sampled the four-dimensional space of activation patterns, all of which produce the same tip force but vary in stiffness and stability (*f_y_* = 9.1 N, *f_x_* = –8.0N). Using the nondimensional variables *ηb/k*_min_ and (*k*_joint_ – *k*_min_)/*k*_min_ for stability and stiffness, respectively, the 4D space of activations collapses into a family of 1D curves that are parametrized by the damping value. Near the origin, the 1D stability-stiffness curves merge into a universal line with slope = –1 according to the asymptotic relation (3). **b,** The unstable optimal activation 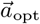 (**inset**) that maximizes tip force, and **c,** the marginally stable pattern ‘2’ are linearly scaled to vary the tip force. The joint stiffness *k*_joint_ and the minimum stiffness *k*_min_ also scale linearly, thus preserving the stability properties of the original activation pattern. (**inset c,**) Maximally scaled up version of pattern ‘2’. Posture for all plots: (30°, 30°, 10°).

We sampled the null-space using 6× 100 million Monte Carlo simulations at 6 different damping values *b* (supplement §S1.5), and show the results on a nondimensional stability-stiffness space (figure 5a). The finger was critically or over damped, like past measurements (Hajian and Howe, 1997). The nondimensional variables are found from asymptotic analysis of the eigenvalue equation (S1.11) near the point of marginal stability when *k*_joint_ = *k*_min_. The minimal stiffness for stability is given by *k*_min_ = *f*_0_*ℓ*, where *f*_0_ is the tip force magnitude and *ℓ* is a posture-dependent length scale (supplement §S1.1). As *k*_joint_ nears *k*_min_, the stability-dominating eigenvalue *η* that has the largest real part is asymptotically given by,

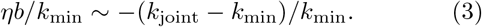

This trade-off between stability *ηb/k*_min_ and stiffness (*k*_joint_ – *k*_min_)/*k*_min_ is a universal (asymptotic) relationship that is independent of the finger’s mass, and accounts for differences in damping, posture, force magnitude, or force direction.

Within the null-space are stable patterns like ‘3’ (supplement §1.5, movie 2) and marginally stable patterns like ‘2’ with stable and unstable patterns on either side of it. Importantly, as the co-contraction decreases and the finger approaches marginal stability, the nondimensional stability-stiffness curves collapse onto a universal line with slope = –1 given by equation (3). Thus, more co-contraction improves stability but also makes the finger stiffer.

We used the model to also examine stability when a specific activation pattern is linearly scaled, in turn linearly scaling the tip force. This is motivated by simultaneous intramuscular EMG recordings from all seven muscles in humans (Valero-Cuevas, 2000; Valero-Cuevas *et al*., 2009; Venkadesan and Valero-Cuevas, 2008), which showed that people exert sub-maximal forces by linearly scaling the activation pattern that they used to stably exert maximal force. We found that the stability characteristics are preserved by linearly scaling the activation pattern because the governing equations (S1.10) and (S1.12) are linear (supplement §S1). So, producing sub-maximal forces by scaling the maximal force pattern 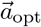 does not help stability and the finger remains unstable (figure 5b). However, for the marginally stable activation pattern ‘2’ that produces sub-maximal force, the marginal stability is preserved whether the activations are scaled up or down (figure 5c).

## III. DISCUSSION

We have shown that maximum exertion of force is limited by stability than muscular capacity, and people restrict how hard they push because the finger would otherwise buckle. Neural feedback control alone cannot help because the buckling instability is too fast relative to sensorimotor latencies during maximal voluntary effort. So people rely on the stiffness arising from muscle contraction. Although the short-range stiffness of muscle is proportional to force, it does not automatically guarantee stability. Only select combinations of muscle contraction and co-contraction patterns can help stiffen and stabilize the finger. Indeed, people are significantly co-contracted when producing fingertip forces, likely for stability. Our co-contraction analysis from EMG measurements and the finger model shows that stiffness due to muscle-induced co-contraction is a viable stabilizing strategy only at sub-maximal forces. That is why, even when instructed to maximize force, people apply lesser force than the capacity of their muscles.

Muscle’s stiffness and open-loop stability of contacts are important not only for maximal force production but also for precision grips using lighter forces. We have shown that if a specific activation pattern lends stability at maximal force, linearly scaling it down to apply lighter forces will also maintain stability. Past experimental measurements lend support to the idea that people rely on the preservation of stability by linear scaling. When instructed to vary their fingertip force magnitude, people linearly scaled the muscle activation pattern that they used for maximal stable voluntary force (Valero-Cuevas, 2000; Valero-Cuevas *et al*., 2009; Venkadesan and Valero-Cuevas, 2008). We have shown that stability at maximal voluntary force is because of muscle-induced stiffness and not feedback control. When people use scaled versions of the stable maximal voluntary pattern, the finger would continue to remain stable without the need for feedback control. So, we infer that people rely on muscle’s shortrange stiffness for stability when using precisions grips with light forces. Thus, open-loop stability may be a key part of human hand dexterity.

Although neural feedback control is not a viable stabilizing strategy to prevent buckling at maximal force, it might still be effective at lighter forces. This is because the real part of the eigenvalue that governs the rate of growth of instabilities could be smaller at lighter forces. For a pattern 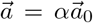 that scales an unstable pattern 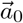 by a factor *α* < 1, the original eigenvalue *η*_0_ becomes *η* = *αη*_0_(*b*_0_/*b*), where *b*_0_ and *b* are the damping associated with the original and scaled patterns, respectively. How damping scales with muscle contraction is presently unclear, but it is hypothesized to scale nonlinearly with activation (Hajian and Howe, 1997; Nguyen *et al*., 2018). So, the eigenvalue could vary with the scaling factor α and the unstable growth of the finger’s posture may be slower at lower activations. A high force that was outside the ability of feedback control to stabilize could thus become stabilizable at lower activation levels. But stability is still a key objective for selecting coordination patterns that allow feedback to augment the role of muscle’s stiffness. Future studies are needed to test whether neural feedback is used in this manner, or whether people rely mostly on open-loop stability based on muscle’s stiffness even when applying light fingertip forces.

In contrast to our finding that neural feedback alone cannot stabilize the finger’s internal degrees of freedom at maximal force, previous studies on endpoint force and position control find that neural feedback is important (Chib *et al*., 2009; Doemges and Rack, 1992; Hu *et al*., 2017; Mugge *et al*., 2009). One difference from our study is that these past studies investigated low forces, where feedback may prove effective. Furthermore, our results do not show that neural feedback is absent, and instead only shows that neural feedback cannot be effective at maximal force. For example, stretch reflexes will exist at maximal force and would likely be elicited by the buckling event. But the rapidity of the event would render the reflexes unable to prevent the finger from buckling. At lower forces, these reflexes and other slower sensorimotor feedback may indeed complement the role of muscle co-contraction and whether that is the case remains to be seen through future studies that may use sensory blocks or other means to investigate the quetion. Nevertheless, the fact that the subjects knew how far to push in order to not buckle indicates that trial-to-trial feedback and learning were used to limit the maximal force and to select appropriate co-contraction strategies that balance the trade-off between force maximization and stability maintenance.

More stiffness is better for postural stability, but has implications for everyday hand usage. Although people pre-shape their hand to match the object about to be gripped (Jeannerod *et al*., 1995), the fingers need to further deform upon contact to adapt to the object’s precise geometry (Santello and Soechting, 1998). Compliance is essential for the fingers to deform and adapt the grasp to the object’s geometry (Erdmann and Mason, 1988; Kao *et al*., 1997; Kazemi *et al*., 2012; Mason, 1981), and is also needed to avoid the well-known instabilities associated with tip force control (Hogan and Buerger, 2018; Whitney, 1987). But the compliance for adaptive grasping has to be traded-off against stiffness for postural stability. More work is needed to understand this trade-off, but the augmentation of open-loop stability by neural feedback at light forces may help manage the trade-off. The universal stability-stiffness curve shows how some patterns may be weakly unstable and allow the finger to be more compliant than strictly enforcing open-loop stability. Thus, our findings on the stability-stiffness trade-off may underlie the selection of strategies for stable yet compliant grasps. Open questions also remain on how such strategic muscle co-contraction is acquired through experiential learning, and how these patterns are related to the vigorously debated neuromuscular synergies (Santello *et al*., 2016; Takei *et al*., 2017; Weiss and Flanders, 2004). Nevertheless, the generality of our results imply that muscle’s role in open-loop stabilization must be considered.

Our results and analyses present some possible generalizations and implications to other areas. Simulation studies have found that contact-induced postural instabilities could also occur in the legs of standing cats and humans (Bunderson *et al*., 2008; De Groote *et al*., 2017). Because inertia is not involved in the universal nondimensional stability-stiffness curve at the margin of stability, our results are applicable to multi-link chains of diverse length scales. As animal limbs typically have more muscles than kinematic degrees of freedom, joint stiffness can be controlled independent of the torques and therefore strategies may exist to use co-contraction to maintain stability. In robotic limbs also, stiffness and torques can be independently controlled because variable impedance actuators are increasingly prevalent (Vanderborght *et al*., 2013). Therefore, our results apply broadly across animals and machines for achieving compliant, adaptive, and stable contacts. Furthermore, our results are also applicable when the condition of a rigid tip contact is relaxed to include a compliant contact, such as during multi-fingered grasps. In force control, contact compliance is beneficial for stability. But the postural instability relies on the tip force magnitude alone, and importantly, the proportionality between the minimum joint stiffness for stability and tip force would remain. What compliant contact, such as in multi-fingered grasps may help with is to reduce the severity or remove the trade-off between compliance for force control versys stiffness for postural stability. Finally, our results have implications for data normalization methods and the use of maximum voluntary contraction (MVC) measurements in the clinical functional testing, and in neuromechanical and biomechanical studies (Burden, 2010; Halaki and Ginn, 2012). Such measurements should either consider externally stabilizing the limb in question or find means to delineate the role of stability versus muscular capacity in maximal EMG and force production.

## IV. METHODS

We conducted two separate experiments with the voluntary participation of right-handed adults, who had no history of hand injuries or impairments: (**i**) experiment 1 to record finger buckling: n=10, 7M, 3F, age 24-47 years, and (**ii**) experiment 2 to test maximal force with splints: n=39, 26M, 13F, age 18-47 years. Subjects studied the consent form and the experimenter discussed potential risks of the study and their option to withdraw from the study at any time. The experiment was performed after the subject provided informed consent and in accordance with the relevant guidelines and regulations. The detailed procedures for seeking informed consent were approved by Yale University’s IRB (HIC# 2000029475). A similar procedure was followed for the study conducted in India, with approval from the Institute Ethics Committee (Human Studies) of the National Centre for Biological Sciences, Bengaluru, India. One subject from experiment 1 was excluded because they never exhibited buckling events, which was the goal of the study. One subject from experiment 2 was excluded because their finger repeatedly buckled and did not yield reliable data.

### A. Experiment 1: Buckling timescale

Subjects maintained a flexed index finger posture of their choosing, with the fingertip pushing on a horizontal steel plate (figure 1). They were asked to maximize the distal fingertip force while maintaining a flexed finger posture with no regard for postural stability. Circular 3mm reflective markers were attached to the radial aspect of the metacarpophalangeal (MCP), proximal inter-phalangeal (PIP) and distal interphalangeal (DIP) joints, the fingertip, and the second metacarpal’s proximal end. A high-speed camera (Photron FASTCAM Mini AX100, MEC, Westfield, IN) recorded the instances of buckling in the lateral (radial) view of the index finger at 4500 fps for Subject 1 and 5000 fps for rest of the subjects. Trials were separated by 2 minutes to reduce fatigue.

We estimated the change in the joint angles Δ*θ_x_* where *x* is either DIP, PIP, or MCP from the videos using custom software. The time-history Δ*θ_x_*(*t*) was used once the angle increased past 2°, and until the subject-dependent end of buckling (supplement table S3). To estimate the time-constant *τ* for the hypothesized exponential growth Δ*θ_x_*(*t*) = Δ*θ*_0_*e^t/τ^*, we performed a linear regression of logΔ*θ_x_*(*t*) versus *t* using the middle half of the data to avoid end-effects associated with the log-transform. The slope of the semi-log plot is equal to 1/*τ* (figure 1, inset). The estimated time-constants and R^2^ of the fits are reported for the thirty-three trials where the finger buckled (supplement table S3). Matlab (version 9.8.0.1323502, Natick, MA) was used for the image and regression analyses.

### B. Experiment 2: Splinted finger

The second experiment tested the model’s predictions by measuring the change in maximal voluntary force when the stability of the finger was altered by externally stiffening it.

#### Experimental apparatus

Subjects wrapped their thumb and unused fingers of the right hand around a ground-mounted non-slip handle and pushed on a horizontal steel plate with their index finger. The handle was adjusted so that the MCP, PIP, and DIP joints were at 30°, 30°, and 10° flexion, respectively (figure 3a). Using established methods (Valero-Cuevas *et al*., 2009, 1998; Venkadesan and Valero-Cuevas, 2008), the fingertip was covered by a subject-specific custom-molded thermoplastic thimble (Türbocast^®^, T-Tape Company, The Netherlands) and fixed using Vetrap bandage (3M, Maplewood, Minnesota) to yield a well-defined contact point and consistent friction.

To stiffen the finger, we attached subject-specific thermoplastic 2-joint and 1-joint splints to the dorsal face of the index finger using Vetrap bandage. The splints were molded to each subject’s finger at the posture (30°, 30°, 10°). The 2-joint splint covered MCP, PIP and DIP joint and stiffened the PIP and DIP joints, but the 1-joint splint covered only the MCP and PIP joints, and stiffened the PIP joint.

#### Experimental protocol

The subjects were asked to try 2-6 times to apply the greatest vertical force they could during the measurement window without letting the finger buckle or the tip slip on the surface, with at least 2 minutes rest between tries. Three finger conditions were tested: free, 2-joint splint (2J), and 1-joint splint (1J). For 9 subjects (set A) we tested free and 2J fingers with a 15s measurement window. For 14 subjects (set B), the free, 1J, and 2J fingers were measured using a 20 s window. For 16 subjects (set C), the free, 1J, and 2J fingers were measured using a 15 s window. To control for motor learning, the order was randomized in set A (free before splint for 7 subjects) and set C (free before splint for 7 subjects). To control for fatigue, the free finger was always first in set B. Subjects were acclimatized to the splint by handling objects and lightly pushing on surfaces before measurement. The vertical fingertip force was displayed as a live trace on a monitor. The fingertip never slipped, but trials where the free finger buckled were excluded.

#### Data recording

Fingertip force was recorded at 2000 Hz by rigidly fixing a six-axis load cell (model 45E15A4-M63J-EF, JR3 Inc., Woodland, CA) between the steel plate and a rigid bench. Surface-EMG was acquired at 2000 Hz using wireless electrodes (Trigno Wireless EMG System, Delsys Inc., Natick, MA). We palpated the ventral side of the forearm when the subject resisted forces on the index finger to identify the two extrinsic flexors, *flexor digitorum profundus* (FDP) and *flexor digitorum superficialis* (FDS), and attached the electrodes to the skin over the muscle belly using hypoallergenic double-sided tape and Vetrap bandage. For Set C, additional EMG was recorded from one extensor, *extensor digitorum communis* (EDC). We verified the electrode placement by asking subjects to push the experimenter’s hand using their index finger while observing the EMG traces.

#### Signal processing

EMG recordings were band-stop filtered in the range 48 – 52 Hz and 98 – 102 Hz with zero phase distortion to remove electrical noise for Set A and B (India uses 50 Hz AC supply), and in the range 58 – 62 Hz and 118 – 122 Hz for Set C (USA uses 60Hz AC supply). We then high pass filtered at 20 Hz to remove movement artifacts, full-wave rectified and passed through a fourth-order Butterworth filter with a time constant of 0.23 s to adjust for the muscle’s excitation-contraction dynamics (Valero-Cuevas *et al*., 2009).

The force and the processed EMG were moving average filtered with a 1-second window (figure 3b) to find the maximal voluntary force F_max_ and the EMG at that time. The fingertip force vector across trials was oriented 5.0±2.8deg (mean±SD) from the vertical and we verified that the results were not sensitive to the moving average window size (figure S2). We normalized the maximal voluntary force F_max_ of each trial by the maximal forces from all the trials of that subject to obtain a normalized force measure *f*_max_. For each subject, we normalized the EMG recordings of each muscle with the moving average filtered maximal activity of the corresponding muscles. A PCSA-weighted average of the smoothed and normalized flexor EMG signals was used to find EMG_flexors_.

#### Statistical analysis

We report the mean±SE of the change in *f*_max_ and EMG_flexors_, calculated as the difference between the splinted and the free conditions, for the 2J and 1J splint to assess whether the force and EMG significantly increased with the splint. Additionally, descriptive statistics for F_max_ and EMG_flexors_, and the change in normalized force *f*_max_ and EMG_flexors_ are in the supplement (supplement table S4 and S7).

Two one-way mixed-model Type III ANOVAs using Satterthwaite’s method tested the effect of splint (free, 1J, 2J) and order of splint conditions on *f*_max_ and EMG_flexors_, with subject as a random factor. Tukey contrasts for multiple comparisons of means were used and Bonferroni-Holm method were applied to find the adjusted p-values (supplement table S6). Two linear regressions, one for each splint type, modeled the relationship between change in *f*_max_ (dependent) and change in EMG_flexors_ (explanatory) (supplement table S8).

For 16 subjects in Set C, we measured EMG_EDC_ along with EMG_FDP_ and EMG_FDP_. An ANOVA tested the effect of finger condition on the ratio of EMG_EDC_ to EMG_flexors_. Three ANOVAs tested the effect of finger condition on the EMG to normalized force ratios for FDS, FDP, and EDC. Bonferroni-Holm method were applied on the multiple comparison of means to find the adjusted p-values (supplement table S10 and S11). Using the difference between the linearly scaled splinted EMG_EDC_ for producing the same normalized maximal force as the free finger and the EMG_EDC_ for the free condition, we calculated the change in co-contraction of EDC (Δcc_EDC_) between the splinted and the free condition (figure 4a). Three linear regressions tested the effect of change in co-contraction Δcc_EDC_ on the change in normalized fingertip force Δ*f*_max_, the effect of free finger’s baseline force F_free_ (N) on the change in co-contraction Δcc_EDC_, and the effect of free finger’s baseline force F_free_ (N) on the change in fingertip force ΔF_max_ (N), respectively.

We verified statistical assumptions of normality and equivariance (supplement figure S3, S4, and S7). Significance level for all statistical tests was *a priori* set to 0.05. RStudio (version 1.1.463, RStudio Team, 2016) was used for the statistical tests. The complete dataset is provided as supplement files.

## ACKNOWLEDGMENTS

P. Paoletti for discussions on constrained dynamics. Reviewers for useful comments. Funding from the Wellcome Trust DBT Alliance to M.V., and Yale’s Integrated Graduate Program in Physical and Engineering Biology, the Pierre W. Hoge Foundation Fund, and the Alpheus B. Stickney Scholarship Fund to N.S. National Centre for Biological Sciences, India, for supporting N.S. during the experiments.

## Author contributions

M.V. conceived the project, N.S. performed the experiments, analysis and simulations, N.S. and M.V. jointly interpreted the results, and wrote the paper.

## REFERENCES

Akella, P., and M. Cutkosky (1989), in Robotics and Automation, 1989. Proceedings., 1989 IEEE International Conference on (IEEE) pp. 764–769.

Bicchi, A., and V. Kumar (2000), in Proceedings 2000 ICRA. Millennium Conference. IEEE International Conference on Robotics and Automation. Symposia Proceedings (Cat. No. 00CH37065), Vol. 1 (IEEE) pp. 348–353.

Bunderson, N. E., T. J. Burkholder, and L. H. Ting (2008), Journal of biomechanics 41 (7), 1537.

Burden, A. (2010), Journal of electromyography and kinesiology 20 (6), 1023.

Burdet, E., R. Osu, D. W. Franklin, T. E. Milner, and M. Kawato (2001), Nature 414 (6862), 446.

Chib, V. S., M. A. Krutky, K. M. Lynch, and F. A. Mussa-Ivaldi (2009), Journal of Neuroscience 29 (12), 3939.

Cui, L., E. J. Perreault, H. Maas, and T. Sandercock (2008), Journal of Biomechanics 41 (9), 1945.

De Groote, F., J. L. Allen, and L. H. Ting (2017), Journal of biomechanics 55, 71.

Doemges, F., and P. Rack (1992), The Journal of Physiology 447 (1), 575.

Erdmann, M. A., and M. T. Mason (1988), IEEE Journal on Robotics and Automation 4 (4), 369.

Hajian, A. Z., and R. D. Howe (1997), Journal of Biomechanical Engineering 119 (1), 109.

Halaki, M., and K. Ginn (2012), Computational intelligence in electromyography analysis-a perspective on current applications and future challenges, 175.

Hanafusa, H., and H. Asada (1977), IFAC Proceedings Volumes 10 (11), 127.

Hogan, N., and S. P. Buerger (2018), “Impedance and interaction control,” in Robotics and automation handbook, edited by T. R. Kurfess, Chap. 19 (CRC Press) pp. 375–398.

Hu, X., D. Ludvig, W. M. Murray, and E. J. Perreault (2017), Scientific reports 7 (1), 1.

Jeannerod, M., M. A. Arbib, G. Rizzolatti, and H. Sakata (1995), Trends Neurosci 18 (7), 314.

Johansson, R. S., and I. Birznieks (2004), Nature Neuroscience 7 (2), 170.

Kao, I., M. R. Cutkosky, and R. S. Johansson (1997), IEEE Transactions on Robotics and Automation 13 (4), 557.

Karakostis, F. A., G. Hotz, V. Tourloukis, and K. Harvati (2018), Science Advances 4 (9), eaat2369.

Kazemi, M., J.-S. Valois, J. A. Bagnell, and N. Pollard (2012), Robotics: Science and Systems (RSS)(MIT Press, Sydney, Australia, 2012), 177.

Kivell, T. L. (2015), Philosophical Transactions of the Royal Society B: Biological Sciences 370 (1682), 20150105.

Klimchik, A., D. Chablat, and A. Pashkevich (2015), European Journal of Mechanics-A/Solids 51, 193.

Marco, R. A., N. A. Sharkey, T. S. Smith, and A. G. Zissimos (1998), JBJS 80 (7), 1012.

Marzke, M. W. (1997), American Journal of Physical Anthropology 102 (1), 91.

Marzke, M. W. (2013), Philosophical Transactions of the Royal Society B: Biological Sciences 368 (1630), 20120414.

Marzke, M. W., and R. F. Marzke (2000), Journal of Anatomy 197, 121.

Mason, M. T. (1981), IEEE Transactions on Systems, Man, and Cybernetics 11 (6), 418.

Mugge, W., J. Schuurmans, A. C. Schouten, and F. C. van der Helm (2009), Journal of Neuroscience 29 (17), 5476.

Murray, R. M., Z. Li, and S. S. Sastry (2017), A mathematical introduction to robotic manipulation (CRC press).

Napier, J. R. (1956), Bone & Joint Journal 38 (4), 902.

Nguyen, K. D., N. Sharma, and M. Venkadesan (2018), Frontiers in Robotics and AI 5, 69.

Okamura, A. M., N. Smaby, and M. R. Cutkosky (2000), in Proceedings 2000 ICRA. Millennium Conference. IEEE International Conference on Robotics and Automation. Symposia Proceedings (Cat. No.00CH37065), Vol. 1, pp. 255–262 vol.1.

Peterka, R. (2002), Journal of neurophysiology 88 (3), 1097.

Rack, P. M., and D. Westbury (1974), The Journal of Physiology 240 (2), 331.

Rancourt, D., and N. Hogan (2001), Journal of Motor Behavior 33 (2), 193.

Santello, M., M. Bianchi, M. Gabiccini, E. Ricciardi, G. Salvietti, D. Prattichizzo, M. Ernst, A. Moscatelli, H. Jorntell, A. M. L. Kappers, K. Kyriakopoulos, A. A. Schaeffer, C. Castellini, and A. Bicchi (2016), Phys Life Rev 17, 54.

Santello, M., and J. F. Soechting (1998), J Neurophysiol 79 (3), 1307.

Schweizer, A. (2001), Journal of biomechanics 34 (2), 217.

Selen, L. P., D. W. Franklin, and D. M. Wolpert (2009), The Journal of Neuroscience 29 (40), 12606.

Takei, T., J. Confais, S. Tomatsu, T. Oya, and K. Seki (2017), Proc Natl Acad Sci U S A 114 (32), 8643.

Valero-Cuevas, F. J. (2000), Journal of Neurophysiology 83 (3), 1469.

Valero-Cuevas, F. J., M. Venkadesan, and E. Todorov (2009), Journal of Neurophysiology 102 (1), 59.

Valero-Cuevas, F. J., F. E. Zajac, and C. G. Burgar (1998), Journal of Biomechanics 31 (8), 693.

Vanderborght, B., A. Albu-Schaeffer, A. Bicchi, E. Burdet, D.G. Caldwell, R. Carloni, M. Catalano, O. Eiberger, W. Friedl, G. Ganesh, M. Garabini, M. Grebenstein, G. Grioli, S. Haddadin, H. Hoppner, A. Jafari, M. Laffranchi, D. Lefeber, F. Petit, S. Stramigioli, N. Tsagarakis, M. Van Damme, R. Van Ham, L. C. Visser, and S. Wolf (2013), Robotics and Autonomous Systems 61 (12), 1601.

Venkadesan, M., and F. J. Valero-Cuevas (2008), The Journal of Neuroscience 28 (6), 1366.

Venkadesan, M., and F. J. Valero-Cuevas (2009), Philosophical Transactions of the Royal Society of London A: Mathematical, Physical and Engineering Sciences 367 (1891), 1163.

Vigouroux, L., F. Quaine, A. Labarre-Vila, and F. Moutet (2006), Journal of biomechanics 39 (14), 2583.

Weiss, E. J., and M. Flanders (2004), Journal of neurophysiology 92 (1), 523.

Whitney, D. (1987), The International Journal of Robotics Research 6 (1), 3.

