## Supplement for "Finger stability in precision grips"

#### This supplement includes:

- Movie 1: high-speed video of a buckling finger
- Movie 2: numerical finger simulations
- Dataset 1: buckling experiments
- Dataset 2: splint experiments
- Supplementary notes
  - S1: mathematical modeling
  - S2: experimental methods and statistics
  - Figures S1 to S7
  - Tables S1 to S13
  - References for SI citations

**Movie 1: High-speed video recording of the buckling instability in the index finger.** The subject maintained a self-selected finger posture and pushed as hard as they could on a steel plate. The change in distal interphalangeal (DIP) joint angle $\Delta\theta_{\mathrm{DIP}}$ once it increased past 2° and until the subject-dependent end of buckling was used to fit an exponential and estimate the time-constant of the unstable growth. For this trial, the time-constant was 20 ms with $\mathrm{R}^2 = 0.998$ for the exponential fit. The video was recorded at 5000 fps.

**Movie 2: Animation of the perturbation response for a muscle-driven index finger model.** This movie shows the response of an anatomically detailed model of the index finger in the posture $(30°, 30°, 10°)$ when subjected to finite perturbations. The animation on the left uses an activation pattern $\vec{a}_{\mathrm{opt}}$ that maximizes the vertical fingertip force ($f_x = -8$ N $f_y = 43$ N). The one on the right produces a sub-maximal force $f_x = -8$ N, $f_y = 9$ N, which uses the muscle activation pattern '3' (main text figure 5). The equilibrium posture $(30°, 30°, 10°)$ is perturbed to $\vec{\theta} = (29.98°, 30.16°, 9.71°)$ at $t = 0$ and the response is simulated for a total of 100 ms. The finger is unstable with maximal force activation pattern but recovers its posture when using a sub-maximal force activation pattern. The damping constant at each joint is 103 N mm ms.

**Dataset 1: Finger buckling experiments with 9 subjects.** Time-coded index finger joint angles in 33 buckling trials from two subjects. The angles were found using photogrammetry of the video recordings of the buckling finger. The distal interphalangeal (DIP), proximal interphalangeal (PIP) and metacarpophalangeal (MCP) joint angles are in degrees, and time is in seconds. A straight finger is defined as zero for all the angles and flexion is positive. The data include the entire trial and not just when the DIP angle increased beyond 2 degrees. The video was recorded at 4500 fps for subject 1 and 5000 fps for the rest of the subjects.

**Dataset 2: Splinted finger experiments with 39 subjects.** Subject ID, splint condition, maximal voluntary force $f_{\max}$ (N), the MVC-normalized EMG from *flexor digitorum profundus* (FDP), *flexor digitorum superficialis* (FDS), *extensor digitorum communis* during the time window of maximal force, the normalized force, and the order of finger conditions are included in the dataset.

### Supplementary notes
### Finger stability in precision grips

Neelima Sharma[1] and Madhusudhan Venkadesan*[1]

[1]*Department of Mechanical Engineering & Materials Science, Yale University, New Haven, CT, USA*

**References** **21**

*

### S1 Mathematical modeling

#### S1.1 Index finger model

The objective for modeling the index finger is to analyze fingertip force application and stability while holding the posture constant. During the experiments, the index fingertip pushed on a rigid surface without slipping, and the knuckle was held in place by holding onto a dowel. So, like previous studies [1, 2], we modeled the index finger as a planar, actuated three-link chain that is hinged to the ground at the knuckle and the fingertip. Open-loop control torques and mechanical elements such as springs and dampers were included, but feedback control was not considered because we want to examine the open-loop stability of the finger.

**Governing equations:** We derive the governing equations of a constrained three-link chain whose fingertip position $\vec{r}_e$ with respect to the knuckle is constant. The joint angles are $\vec{\theta} \in \mathbb{R}^3$, the inertia matrix is $M(\vec{\theta}) \in \mathbb{R}^{3\times3}$, the torques due to Coriolis and centripetal forces are $\vec{c}(\vec{\theta}, \dot{\vec{\theta}}) \in \mathbb{R}^3$, the posture-dependent endpoint Jacobian matrix is $J(\vec{\theta}) \in \mathbb{R}^{2\times3}$, the fingertip force due to the constraint is $\vec{f}(\vec{\theta}, \dot{\vec{\theta}}) \in \mathbb{R}^2$, the constant open-loop actuation torques are $\vec{\tau}_0 \in \mathbb{R}^3$, the joint stiffness matrix is $K \in \mathbb{R}^{3\times3}$, and the joint damping matrix is $B \in \mathbb{R}^{3\times3}$. The neutral joint angles $\vec{\theta}_0$ for the stiffness term is chosen to be the static posture of choice, and are also used to specify the conditions for static equilibrium and for linearizing the equations.

Omitting the dependence on the joint angles and angular velocities for brevity, the dynamical equations, constraints, and constraint forces are, respectively,

$$M\ddot{\vec{\theta}} + \vec{c} + K(\vec{\theta} - \vec{\theta}_0) + B\dot{\vec{\theta}} + J^T\vec{f} = \vec{\tau}_0, \tag{S1.1a}$$

$$\vec{r}_e = \text{constant, and} \tag{S1.1b}$$

$$\vec{f} = (JM^{-1}J^T)^{-1}(JM^{-1}(\vec{\tau}_0 - \vec{c} - K(\vec{\theta} - \vec{\theta}_0) - B\dot{\vec{\theta}}) + \dot{J}\dot{\vec{\theta}}). \tag{S1.1c}$$

**Conditions for static equilibrium:** We seek the torques $\vec{\tau}_0$ for the finger to be in equilibrium at the posture $\vec{\theta}_0$ while experiencing the fingertip force $\vec{f}_0$. Setting $\vec{\theta} = \vec{\theta}_0$, $\vec{f} = \vec{f}_0$, $\dot{\vec{\theta}} = \ddot{\vec{\theta}} = 0$ in equation (S1.1a), and using $J_0 = J(\vec{\theta}_0)$, we find

$$\vec{\tau}_0 = J_0^T\vec{f}_0. \tag{S1.2a}$$

Note that given any desired posture $\vec{\theta}_0$ and fingertip force $\vec{f}_0$, we can always find torques that satisfy static equilibrium. However, not every torque leads to static equilibrium because the finger still has one net degree of freedom after imposing the fingertip constraints equation (S1.1b). Using the equilibrium condition (S1.2a) and the relationship (S1.1c) between torque and fingertip force, we arrive at the condition for an applied torque $\vec{\tau}_{eq}$ to maintain equilibrium, i.e. $\dot{\vec{\theta}} = \ddot{\vec{\theta}} = 0$, as

$$\vec{\tau}_{eq} = J^T\left(JM^{-1}J^T\right)^{-1}JM^{-1}\vec{\tau}_{eq}, \tag{S1.3a}$$

$$\iff \vec{\tau}_{eq} \in \text{null}\left(I - J^T\left(JM^{-1}J^T\right)^{-1}JM^{-1}\right). \tag{S1.3b}$$

An applied joint torque satisfies equilibrium if and only if it belongs to the null-space of $(I - J^T(JM^{-1}J^T)^{-1}JM^{-1})$. The matrix $J^T(JM^{-1}J^T)^{-1}JM^{-1} \in \mathbb{R}^{3\times3}$ is a projection matrix [3]. Therefore, its eigenvalues are either 1 or 0 and the number of ones defines the dimensionality of the feasible torque space. For the index finger model, there is a one-dimensional null-space.

**Muscle model:** The joint torques $\vec{\tau}_0$ and joint stiffness K arise from muscle contraction. We use the model developed by Valero-Cuevas [1] for modeling force production in muscles of the index finger, and the model developed by Cui et al. [4] for the short-range stiffness of muscle that induces stiffness at the joints. Because we consider a static posture, the force-velocity properties and the stiffness due to the force-length properties of muscle are inconsequential.

The human index finger has seven muscles, and their activation is parameterized by the activation vector $\vec{a} \in \mathbb{R}^7$. The activations $\vec{a}$ map to muscle forces and muscle stiffness, which lead to joint torques and joint stiffnesses, respectively. This mapping depends on the diagonal matrix $F_{iso} \in \mathbb{R}^{7\times 7}$ of maximal isometric force of each muscle, the matrix $R \in \mathbb{R}^{3\times 7}$ of moment arms of each musculotendon about each joint, the diagonal matrix $L_{iso}$ of isometric lengths for each muscle, and the empirical short-range stiffness factor $\gamma$ that was found by Cui et al. [4]. Thus, the joint torques $\vec{\tau}_0$ and joint stiffnesses K are given by,

$$\vec{\tau}_0 = R F_{iso} \vec{a} \tag{S1.4a}$$

$$K = R\left(\mathrm{diag}\left(\gamma F_{iso} L_{iso}^{-1} \vec{a}\right)\right) R^T. \tag{S1.4b}$$

**Linearized equations:** The linearized governing equations (S1.1a) with constant joint torques $\vec{\tau}_0$ and with small changes in the joint angles $\vec{\theta}' = \vec{\theta} - \vec{\theta}_0$ are,

$$M|_{\vec{\theta}_0} \ddot{\vec{\theta}}' + B\dot{\vec{\theta}}' + \left(K + \left.\frac{\partial (J^T \vec{f})}{\partial \vec{\theta}}\right|_{\vec{\theta}_0, \vec{\tau}_0}\right) \vec{\theta}' = 0. \tag{S1.5}$$

The angle changes $\vec{\theta}'$ are subject two constraints (S1.1b), which yields one net degree of freedom $\phi \in \mathbb{R}$. In the linearized setting, this degree of freedom belongs to the null-space of the tip constraints. To derive $\phi$, we linearize the constraint equation (S1.1b) and find its null-space $P \in \mathbb{R}^3$ as,

$$\vec{r}_e = \text{constant} \implies J_0 \vec{\theta}' = 0, \tag{S1.6a}$$

$$J_0 = \left.\frac{\partial \vec{r}_e}{\partial \vec{\theta}}\right|_{\vec{\theta}_0}, \text{ and} \tag{S1.6b}$$

$$P = \mathrm{null}\left(J_0\right). \tag{S1.6c}$$

(S1.6d)

Therefore, in terms of the the null-space P, the reduced degree of freedom is found as a linear combination of the three joint angles according to,

$$\phi = P^T \vec{\theta}'. \tag{S1.7}$$

**Constrained dynamics of the linearized system:** The linearized dynamics of the reduced degree of freedom $\phi$ are obtained by projecting equation (S1.5) onto the null-space of the linearized constraints, which results in a similarity (or coordinate) transform using P. The magnitude of the equilibrium fingertip force $\vec{f}_0$ sets a force scale in the dynamics and we introduce an additional symbol for convenience,

$$f_0 = \|\vec{f}_0\|. \tag{S1.8}$$

The reduced order dynamics for the scalar degree of freedom $\phi$ involve projections of M, B and K, and also a new length scale $\ell$ that relates to the sensitivity of the joint torques to changes in orientation of the fingertip force vector. Thus, by combining equations (S1.5) and (S1.7), we find

$$m\ddot{\phi} + b\dot{\phi} + (k_{\text{joint}} - f_0\ell)\phi = 0, \tag{S1.9}$$

where,

$$m = \mathrm{P}^{\mathrm{T}}\, \mathrm{M}|_{\vec{\theta}_0}\, \mathrm{P}, \tag{S1.10a}$$

$$b = \mathrm{P}^{\mathrm{T}}\mathrm{B}\mathrm{P}, \tag{S1.10b}$$

$$k_{\text{joint}} = \mathrm{P}^{\mathrm{T}}\mathrm{K}\mathrm{P}, \tag{S1.10c}$$

$$\ell = -\frac{1}{f_0}\mathrm{P}^{\mathrm{T}} \left( \left. \frac{\partial(\mathrm{J}^{\mathrm{T}}\vec{f})}{\partial\theta} \right|_{\vec{\theta}_0,\vec{\tau}_0} \right) \mathrm{P}. \tag{S1.10d}$$

#### S1.2 Linear stability analysis

Kinematic chains with more internal degrees of freedom than constraints can buckle under compressive external forces, even when the forces arise due to internal actuation in these mechanisms. To see that, we perform a linear stability analysis of the index finger model that were derived in §S1.1. Recall that the model has one net degree of freedom because there are three internal degrees of freedom and two external constrains on the tip, i.e. an actuated four-bar linkage.

The origin ($\phi = 0, \dot{\phi} = 0$) is the only equilibrium point for the reduced order equation (S1.9), and the eigenvalues of that equilibrium are given by,

$$\eta = \frac{b}{2m}\left(-1 \pm \sqrt{1 - \frac{4m}{b^2}(k_{\text{joint}} - f_0\ell)}\right). \tag{S1.11}$$

The equilibrium is unstable in the sense of Lyapunov when $\mathrm{Re}(\eta) > 0$, i.e. the real part of any one of the eigenvalues is greater than zero. Therefore, the equilibrium is marginally or completely stable in the sense of Lyapunov when,

$$k_{\text{joint}} \geq k_{\text{min}}, \tag{S1.12a}$$

$$k_{\text{min}} = f_0\ell. \tag{S1.12b}$$

Notable properties of the stability criterion include: (**i**) greater magnitude $f_0$ of the fingertip force is more destabilizing, and (**ii**) the condition for stability is independent of damping or inertia, and only depends on the projected stiffness $k_{\text{joint}}$. However, the precise values of the real and imaginary parts of the eigenvalue, when not equal to zero, depend on inertia and damping.

#### S1.3 Optimal activation pattern to maximize fingertip force

We combine equations (S1.1c), (S1.3a), and (S1.4a) to define an optimization problem to find the activation pattern that maximizes the fingertip force while maintaining equilibrium. Following the instructions to the subjects, we choose the vertical component of the fingertip force as the objective function. The optimization problem thus stated is,

$$\begin{aligned} &\max_{\vec{a}} \begin{pmatrix} 0 \\ 1 \end{pmatrix} \cdot \vec{f}, \\ &\text{subject to:} \\ &\vec{f} = (\mathrm{J}\mathrm{M}^{-1}\mathrm{J}^{\mathrm{T}})^{-1}\mathrm{J}\mathrm{M}^{-1}\mathrm{R}\mathrm{T}\vec{a}, \\ &\mathrm{J}^{\mathrm{T}}\vec{f} = \mathrm{R}\mathrm{F}_{\text{iso}}\vec{a}, \\ &0 \leq a_i \leq 1, \text{ for } i = 1, \ldots, 7. \end{aligned} \tag{S1.13}$$

The optimization problem is linear in the objective and constraint functions, and therefore easily solved using standard linear programming solvers. These equations are analogous to the approach of Valero-Cuevas [1], with the difference that we incorporate the equilibrium constraint using a different form of the governing equations. Valero-Cuevas [1] adds a dummy rotational torque at the fingertip to make a square Jacobian that can be inverted to solve the tip force for any given joint torque, and then imposes the constraint that the dummy torque should be zero.

##### S1.4 Numerical analysis of stability

**Parameter values:** The muscle and segment parameters used for analysis of the finger are provided in tables S1 and S2.

**Maximal force:** We found optimal activations that maximize fingertip force by following the procedure of §S1.3. The optimization was performed within Matlab version 9.8.0.1323502 (R2020a, Natick, MA) using CVX, an optimization package for convex programs [5, 6].

The results of the optimization are reported in figure 2 in the main text of the paper. For the posture used in our experiments, $\vec{\theta} = \begin{pmatrix} 30^\circ & 30^\circ & 10^\circ \end{pmatrix}^T$, we found the optimal activation pattern $\vec{a} = \begin{pmatrix} 1 & 0.52 & 0 & 0 & 1 & 1 & 1 \end{pmatrix}^T$, and maximal vertical force $f_{max} = 42.6\,\text{N}$ that is pointed distally. The fingertip force vector also had a horizontal component $f_x = -8.0\,\text{N}$ that is pointed towards the palm.

**Stability at maximal force:** We calculated the minimum stiffness $k_{min}$ that is needed for stability of the maximal force solution and compared it against the joint stiffness $k_{joint}$ that is induced by the muscle activation pattern. For the force-maximizing activation pattern, $k_{min} = 295\,\text{N mm}$ and $k_{joint} = 109\,\text{N mm}$. Therefore, the force-maximizing activation pattern makes the finger posturally unstable (see also §S1.4, movie 2).

The one internal degree of freedom implies a one-parameter family of postures all which preserve the tip constraint. We found the maximal force at 3991 postures that spanned joint angles without hyper-extending any of them. None of those postures were stable as seen from figure S1a that shows that $k_{joint} < k_{min}$ for all of those postures.

**Stability at sub-maximal force:** In addition to the maximal force, we examined activation patterns that produced sub-maximal force in the vertical direction, but with the same horizontal force as the maximal force pattern. We show an illustrative example of such an activation pattern that produces sub-maximal force (see §S1.5), namely, $\vec{a} = \begin{pmatrix} 0.57 & 0.18 & 0.50 & 0.50 & 0 & 0.23 & 0.36 \end{pmatrix}^T$, that applies a sub-maximal vertical fingertip force of $9.1\,\text{N}$ and the same horizontal component as the maximal force, namely $-8.0\,\text{N}$. The finger was posturally stable with this activation because $k_{joint} = 45\,\text{N mm}$ is greater than the minimum stiffness for stability at that force $k_{min} = 39\,\text{N mm}$ (see also §S1.4, movie 2).

**Choice of damping:** For purposes of illustration and visualization of the response to finite perturbations, we chose a diagonal damping matrix B with each joint having a damping constant of $103\,\text{N mm ms}$. The units used reflect the relevant length and time scales for the responses observed in simulations. This damping constant was chosen to illustrate a finger that is weakly under-damped or slightly over-damped in most visualizations. However, the numerical value of the damping has no influence on the stability criterion that is given by equation (S1.12a).

**Response to non-infinitesimal perturbations:** The linearized stability analysis showed the finger's response to infinitesimal perturbations. So, we performed numerical simulations of two test cases using the nonlinear governing equations (S1.1) to study the response to non-infinitesimal perturbations. We chose the activation patterns corresponding to maximal unstable force, and the sub-maximal stable force. We perturbed the equilibrium posture of $\vec{\theta}_0 = (30^\circ, 30^\circ, 10^\circ)$ to

$\vec{\theta} = (29.98°, 30.16°, 9.71°)$ at $t = 0$ and the response was simulated for a total of 100 ms (movie 2). The finger became unstable under the maximal force activation and buckled in a manner similar to our experiments (movie 1). However, the finger was stable and recovered its posture when using the activation pattern that produced sub-maximal force.

##### S1.5 Monte Carlo samples of the activation null-space

We want to investigate the properties of the null-space of the mapping from muscle activations to joint torques. This space defines the set of all activation patterns that can support a specified fingertip force. The steps to numerically chart out the null space are to: (**i**) derive a spanning basis for the null-space, (**ii**) find a particular solution that belongs to the null-space, and (**iii**) sample the space using Monte Carlo simulations.

**Spanning basis for the null-space:** For applying a tip force $\vec{f}$ and keeping the finger in equilibrium, the muscle activation pattern $\vec{a}$ must support the joint torques $\vec{\tau}$ induced by the tip force. The null-space is defined in terms of the manipulator Jacobian J, the muscle force matrix $F_{iso}$, and the moment-arm matrix R, according to,

$$\vec{\tau} = J^T \vec{f} = RF_{iso}\vec{a}, \tag{S1.14a}$$

$$\implies \vec{a}_{null} \in \text{null}(RF_{iso}), \text{ such that} \tag{S1.14b}$$

$$\vec{a} \in \mathbb{R}^7, \ \vec{\tau} \in \mathbb{R}^3, \ \vec{f} \in \mathbb{R}^2, \ RF_{iso} : \mathbb{R}^7 \to \mathbb{R}^3, \text{ and null}(RF_{iso}) = \mathbb{R}^4. \tag{S1.14c}$$

By definition, $RF_{iso}\vec{a}_{null} = 0$. Therefore, if a particular activation $\vec{a}_p$ satisfies the torque equation (S1.14a), then so does $\vec{a}_p + \vec{a}_{null}$. Adding random linear combinations of the basis vectors of the null-space of $RF_{iso}$ to the particular pattern $\vec{a}_p$ yields a random activation pattern that will also satisfy equation (S1.14a). The joint stiffness for each activation is found using equations (S1.4b) and (S1.10c). The eigenvalues are found using equation (S1.11), and depend on the damping constant $b$. For the chosen damping values, the damping coefficient $\zeta = b/(2\sqrt{(k_{joint} - k_{min})m}) \geq 1$ and the finger is critically damped or over damped like in past measurements [7].

**Finding a particular activation pattern:** We used the posture $(30°, 30°, 10°)$ and set the desired sub-maximal tip force to be $f_y = 9.1$ N and $f_x = -8.0$ N. First, we found the bounding activation patterns '1' and '4' that respectively correspond to the minimum and maximum joint stiffness while still belonging to the null space (figure 5a in the main text). To find these extremal patterns, we adapted the linear program in equations (S1.13). We set the objective function as $-k_{joint}$ to find the minimal pattern '1', and as $+k_{joint}$ to find maximal pattern '4'. The average of the minimal and maximal stiffness patterns is used as the particular activation pattern for the Monte Carlo sampling in the next step (pattern '3' in figure 5a in the main text).

**Monte Carlo sampling:** For each damping value chosen, we sampled the null-space by adding 100 million random linear combinations of the basis vectors to the particular pattern '3', while also requiring $0 \leq a_i \leq 1$, for $i = 1, \ldots, 7$. For the illustrative example in the main paper (§IIE), we used the posture $(30°, 30°, 10°)$, a sub-maximal tip force ($f_y = 9.1$ N and $f_x = -8.0$ N), and damping values of $101, 102, 103, 107, 115$, and $136$ N mm ms. A total of 6×100 million Monte Carlo samples were produced. Of these, approximately 6×50 million samples satisfied the bounds that the activations should lie between 0 and 1, and were used to generate the stability-stiffness space in the main text (figure 5).

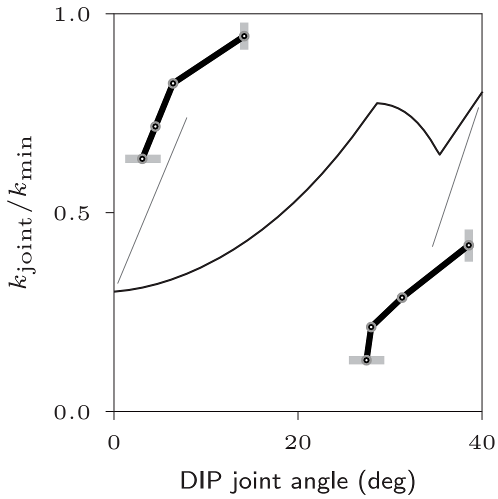

**Fig. S1: Comparison of $k_{joint}$ and $k_{min}$ for multiple postures**. The ratio of the muscle-induced joint stiffness $k_{joint}$ to the minimum stiffness needed for stability $k_{min}$ at a variety of postures. The finger was unstable at all postures.

**Table S1: Muscles used for actuation and their properties.** Seven musculotendon units: *flexor digitorum profundus* (FDP), *flexor digitorum superficialis* (FPS), *extensor indicis proprius* (EIP), *extensor digitorum communis* (EDC), *first lumbrical* (LUM), *first dorsal interosseous* (DI), and *first palmar interosseous* (PI) - actuate the three index finger joints, namely metacarpophalangeal joint (MCP), proximal interphalangeal joint PIP and distal interphalangeal joint (DIP).

| | FDP | FDS | EIP | EDC | LUM | DI | PI |
|---|---|---|---|---|---|---|---|
| Moment arms at finger joints, R (mm) [1, 8] | | | | | | | |
| MCP joint | 12.00 | 13.20 | -7.77 | -7.77 | 7.00 | 2.00 | 4.00 |
| PIP joint | 6.50 | 5.85 | -2.75 | -2.75 | -2.75 | 0 | -2.75 |
| DIP joint | 3.64 | 0 | -1.50 | -1.50 | -1.50 | 0 | -1.50 |
| Muscle physiological properties | | | | | | | |
| PCSA ($cm^2$) [1, 8] | 4.10 | 3.65 | 1.12 | 1.39 | 0.36 | 4.16 | 1.60 |
| Muscle tension (N) [1, 8] | 143.50 | 127.75 | 39.20 | 48.65 | 12.60 | 145.60 | 56.00 |
| Optimal lengths (cm) | 7.5 [9] | 8.4 [9] | 5.9 [9] | 7.0 [9] | 4.9 [10] | 2.9 [11] | 2.9 [12] |

**Table S2: Manipulator properties** are from Venkadesan and Valero-Cuevas, 2009 [2].

| | Proximal phalanx | Intermediate phalanx | Distal phalanx |
|---|---|---|---|
| Lengths (mm) | 50.80 | 25.40 | 19.05 |
| Masses (g) | 8 | 4 | 3 |

### S2 Experimental methods and statistics

#### S2.1 Buckling experiment

**Table S3: Data on the timescale of buckling instability from 9 subjects.** Data are provided for initial and final postures, time constant obtained from the linear regression of $\log(\Delta\theta_x)$ versus time where $x$ is either MCP, PIP, or DIP, and $R^2$ for individual trials.

| Sub | Trial | Joint | $\Delta\theta$ (deg) | $\tau$ (ms) | $R^2$ | Initial posture (deg) (MCP, PIP, DIP) | Final posture (deg) (MCP, PIP, DIP) |
|---|---|---|---|---|---|---|---|
| B | 1 | MCP | 2 to 18 | 55.98 | 0.991 | 8.1, 8.9, 32.5 | -11.8, 17.2, 33.4 |
| | 3 | DIP | 2 to 20 | 52.81 | 0.993 | -38.3, 43.4, 14.5 | -35.3, 48.4, -5.5 |
| | 4 | DIP | 2 to 40 | 37.23 | 0.998 | -12.6, 42.5, 13.0 | -9.7, 62.6, -33.8 |
| | 5 | DIP | 2 to 40 | 77.96 | 0.996 | 9.1, 42.2, 19.9 | 13.8, 60.7, -21.9 |
| | 6 | DIP | 2 to 30 | 42.18 | 0.995 | -8.2, 38.1, 10.1 | -11.9, 58.1, -23.5 |
| H | 6 | DIP | 2 to 30 | 39.45 | 0.995 | -37.1, 51.3, 15.5 | -37.2, 48.8, -13.6 |
| | 8 | DIP | 2 to 23 | 54.80 | 0.900 | -31.2, 54.0, 7.5 | -33.7, 54.5, -14.9 |
| E | 2 | DIP | 2 to 35 | 21.79 | 0.982 | 5.6, 27.8,16.9 | -17.2, 68.1, -19.1 |
| | 4 | MCP | 2 to 20 | 51.17 | 0.984 | 9.2, 28.4, 42.6 | -12.6, 49.4, 31.4 |
| G | 5 | DIP | 2 to 40 | 15.08 | 0.972 | -11.4, 22.9, 28.1 | -14.9, 58.9, -12.3 |
| | 7 | DIP | 2 to 35 | 39.07 | 0.992 | -0.1, 31.2, 35.9 | -0.6, 48.3,-0.4 |
| | 9 | DIP | 2 to 30 | 28.52 | 0.911 | -27.3, 11.9, 22.1 | -27.6, 43.4, -8.6 |
| | 10 | DIP | 2 to 35 | 10.61 | 0.998 | -2.5, 41.3, 40.5 | 10.9, 60.9, -10.9 |
| | 11 | DIP | 2 to 30 | 20.19 | 0.999 | 9.4, 39.9, 29.7 | 8.2, 55.6, -4.7 |
| K | 4 | DIP | 2 to 25 | 43.12 | 0.995 | 29.4, 22.9, 26.4 | 43.9, 24.4, -3.5 |
| | 6 | PIP | 2 to 11 | 55.43 | 0.989 | 35.8, 18.1, 11.8 | 37.5, 5.3, 24.7 |
| | 9 | PIP | 2 to 12 | 17.81 | 0.991 | 37.7, 20.8, 29.5 | 37.0, 3.9, 31.2 |
| M | 7 | DIP | 2 to 25 | 61.29 | 0.996 | -24.2, 39.8, 8.1 | -23.4, 49.9, -19.7 |
| | 8 | DIP | 2 to 23 | 57.90 | 0.993 | -29.5, 34.9, 11.8 | -27.9, 41.7, -12.6 |
| | 9 | DIP | 2 to 25 | 43.49 | 0.983 | -25.9, 37.9, 8.3 | -24.6, 51.0, -17.9 |
| O | 3 | DIP | 2 to 23 | 33.08 | 0.982 | -5.2, 73.4, 13.3 | -6.6, 67.8, 39.2 |
| | 7 | DIP | 2 o 30 | 60.16 | 0.995 | -30.0, 51.8, 16.2 | -29.9, 45.2, 48.9 |
| | 12 | DIP | 2 to 25 | 63.61 | 0.973 | -26.7, 52.9, 30.5 | -30.2, 42.8, 60.8 |
| 1 | 1 | DIP | 2 to 40 | 79.6 | 0.98 | 17.3 , 29.9, -5.3 | 19.4, 46.3, -71.3 |
| | 2 | DIP | 2 to 40 | 21.7 | 0.988 | 6.3, 32.6, 10.5 | -25.1, 69.4, -29.8 |
| | 3 | DIP | 2 to 40 | 14.8 | 0.994 | 8.4, 32.3, -0.8 | -21.1, 74.0, -54.4 |
| | 4 | DIP | 2 to 40 | 46.6 | 0.993 | 15.6, 36.3, -0.7 | 20.8, 55.4, -69.6 |
| 2 | 1 | DIP | 2 to 40 | 12.2 | 0.993 | 56.6, 24.3, 3.0 | 64.5, 37.9, -49.6 |
| | 2 | PIP | 2 to 25 | 42.0 | 0.997 | 55.6, 20.1, 0.8 | 79.9, -7.7, -49.9 |
| | 3 | DIP | 2 to 40 | 27 | 0.972 | 33.1, 18.0, -6.1 | 63.3, -22.4, -54.0 |
| | 4 | DIP | 2 to 40 | 20.4 | 0.998 | -7.7, 35.6, 12.5 | -6.4, 57.0, -46.9 |
| | 5 | DIP | 2 to 40 | 34.2 | 0.985 | -0.2, 36.5, 1.3 | -2.8, 59.9, -55.0 |
| | 6 | DIP | 2 to 40 | 11.1 | 0.997 | -7.1, 41.8, 13.8 | -5.0, 60.7, -51.1 |

#### S2.2 Splint experiment

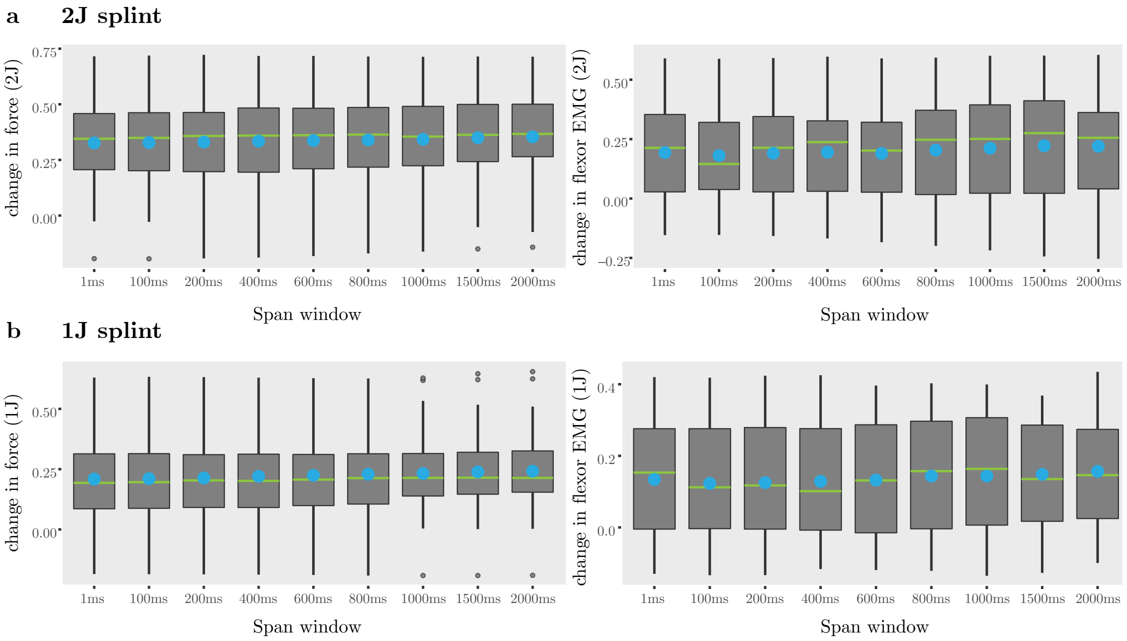

**Fig. S2: Effect of the moving average window size on $\Delta\,f_{max}$ and $\Delta\,EMG_{flexors}$.** Boxplots show the effect of window size ranging from 1 ms to 2000 ms used for moving average filtering. The horizontal green lines in the boxplots identify the median, the blue dots are the mean, the vertical span of the box represents the inter-quartile range, and the whiskers show 1.5 times the interquartile range.

**Table S4: Descriptive statistics of** $F_{max}$ **and** $\mathbf{EMG}_{flexors}$ for no-splint (n=39), 1-joint splint (n=29), and 2-joint (n=38) splint conditions. Mean, standard deviation (SD), $25^{th}$ percentile, median, and $75^{th}$ percentile are reported.

| Splint | Measure | Mean | SD | $25^{th}$ percentile | Median | $75^{th}$ percentile |
|---|---|---|---|---|---|---|
| No splint | $F_{max}$ (N) | 31.51 | 14.93 | 20.88 | 28.86 | 39.26 |
| | $EMG_{flexors}$ | 0.469 | 0.199 | 0.336 | 0.456 | 0.617 |
| 1-joint | $F_{max}$ (N) | 42.46 | 17.23 | 29.49 | 39.99 | 53.67 |
| | $EMG_{flexors}$ | 0.623 | 0.164 | 0.547 | 0.646 | 0.727 |
| 2-joint | $F_{max}$ (N) | 50.96 | 18.87 | 36.55 | 47.69 | 63.85 |
| | $EMG_{flexors}$ | 0.668 | 0.186 | 0.587 | 0.709 | 0.789 |

**Table S5: ANOVA for the effect of splint and order of conditions on $f_{max}$ and $EMG_{flexors}$.** Type III Analysis of Variance with Satterthwaite's method for the effect of splint, order, and the interaction between splint and order on $f_{max}$ and $EMG_{flexors}$ reveals that the effect of splint on $f_{max}$ and $EMG_{flexors}$ is significant whereas the effect of order and the interaction between splint and order is not significant.

| Dependent variable | Condition | Sum sq | Mean sq | Num DF | Den DF | F-value | p-value |
|---|---|---|---|---|---|---|---|
| $f_{max}$ | Splint | 5.86 | 2.93 | 2 | 263.35 | 144.57 | <2e-16 |
| | Order | 0.04 | 0.04 | 1 | 217.97 | 2.13 | 0.15 |
| | Splint:Order | 0.09 | 0.05 | 2 | 67.99 | 2.29 | 0.11 |
| $EMG_{flexors}$ | Splint | 1.777 | 0.889 | 2 | 262.762 | 40.224 | 6e-16 |
| | Order | 0.010 | 0.010 | 1 | 189.727 | 0.459 | 0.49 |
| | Splint:Order | 0.104 | 0.052 | 2 | 69.481 | 2.363 | 0.10 |

**Table S6: Post hoc test for the effect of splint on $f_{max}$ and $EMG_{flexors}$.** Post hoc multiple comparison of means was performed by using Tukey contrasts and adjusted p-values were obtained by applying the Bonferroni-Holm correction.

| Dependent variable | Null hypothesis | Estimate | SE | Z-value | p-value |
|---|---|---|---|---|---|
| $f_{max}$ (N) | 1J - Free == 0 | 0.20 | 0.04 | 4.77 | 6e-6 |
| | 2J - Free == 0 | 0.28 | 0.04 | 6.87 | 2e-11 |
| | 2J - 1J == 0 | 0.08 | 0.04 | 2.19 | 0.09 |
| $EMG_{flexors}$ | 1J - Free == 0 | 0.120 | 0.049 | 2.465 | 0.04 |
| | 2J - Free == 0 | 0.116 | 0.047 | 2.450 | 0.04 |
| | 2J - 1J == 0 | -0.005 | 0.038 | -0.120 | 1 |

**Table S7: Descriptive statistics of $\Delta f_{max}$ and $\Delta EMG_{flexors}$.** Mean, standard error of the mean (SE), $25^{th}$ percentile, median, and $75^{th}$ percentile are reported.

| Splint | Measure | Mean | SE | $25^{th}$ percentile | Median | $75^{th}$ percentile |
|---|---|---|---|---|---|---|
| 2-joint splint | $\Delta f_{max}$ | 0.34 | 0.03 | 0.22 | 0.35 | 0.49 |
| (n=38) | $\Delta EMG_{flexors}$ | 0.212 | 0.034 | 0.022 | 0.251 | 0.394 |
| 1-joint splint | $\Delta f_{max}$ | 0.23 | 0.03 | 0.13 | 0.21 | 0.31 |
| (n=29) | $\Delta EMG_{flexors}$ | 0.143 | 0.028 | 0.006 | 0.164 | 0.307 |

**Table S8: Relationship between $\Delta\,f_{max}$ and $\Delta\,\mathbf{EMG}_{flexors}$.** Results from linear regression to model the relationship between $\Delta\,f_{max}$, the dependent variable, and $\Delta\,EMG_{flexors}$, the explanatory variable.

Model: $\Delta f_{max} = \alpha + \beta\,\Delta EMG_{flexors}$

| Splint | Measure | Estimate | SE | t-value | p-value |
|---|---|---|---|---|---|
| 2-joint splint | $\alpha$ | 0.209 | 0.039 | 5.419 | 4e-6 |
| (n=38) | $\beta$ | 0.626 | 0.132 | 4.754 | 3e-5 |
| | $R^2$=0.39 | | | | |
| 1-joint splint | $\alpha$ | 0.116 | 0.035 | 3.363 | 2e-3 |
| (n=29) | $\beta$ | 0.809 | 0.166 | 4.872 | 4e-5 |
| | $R^2$=0.47 | | | | |

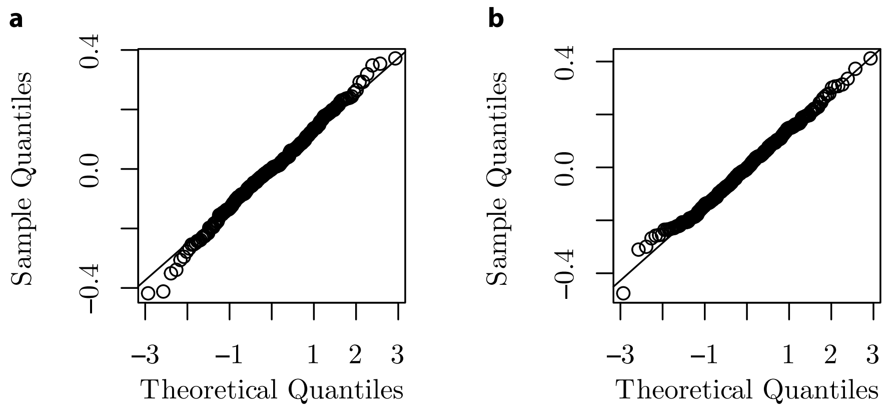

**Fig. S3: Normal probability plot of the residuals obtained from the linear model used in table S5** show that the residuals are normally distributed. Residuals from the linear model that tests the effect of splint, order of splint conditions, and the interaction between order and splint conditions on (**a**,) $\Delta f_{max}$, and (**b**,) $\Delta EMG_{flexors}$.

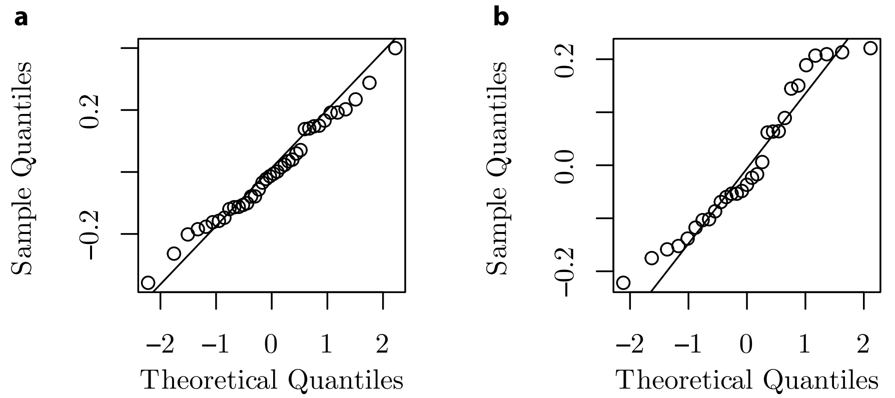

**Fig. S4: Normal probability plot of the residuals for the linear model between** $\Delta f_{max}$ **and** $\Delta \mathbf{EMG}_{flexors}$ (in table S8) for the (**a**,) 2-joint splint and the (**b**,) 1-joint splint.

**Table S9: Relationship between $\Delta F_{max}$ (N) and free finger's baseline force $F_{free}$ (N)**. Results from linear regression to model the relationship between $\Delta F_{max}$ (N), the dependent variable, and $F_{free}$ (N) of the free finger, the explanatory variable.

Model: $\Delta F_{max} = \alpha + \beta F_{free}$

| | | Estimate | Standard Error | t-value | p-value |
|---|---|---|---|---|---|
| 2-joint splint | $\alpha$ (N) | 28.67 | 6.98 | 4.11 | 2e-4 |
| (n=38) | $\beta$ | -0.25 | 0.21 | -1.22 | 0.23 |
| | $R^2 = 0.04$ | | | | |
| 1-joint splint | $\alpha$ (N) | 18.79 | 5.83 | 3.22 | 3e-3 |
| (n=29) | $\beta$ | -0.20 | 0.17 | -1.16 | 0.26 |
| | $R^2 = 0.05$ | | | | |

#### S2.3 Co-contraction analysis

**Table S10: Post hoc contrasts for the effect of finger condition on** $EMG_{EDC}/EMG_{flexors}$. Post hoc multiple comparison of means was performed using Tukey method and adjusted $p$-values were obtained by applying the Bonferroni-Holm correction.

| Contrast | Estimate | Std. Error | df | t-ratio | p-value |
|---|---|---|---|---|---|
| Free - 1J | 0.418 | 0.179 | 45 | 2.338 | 0.06 |
| Free - 2J | 0.435 | 0.179 | 45 | 2.436 | 0.049 |
| 1J - 2J | 0.018 | 0.179 | 45 | 0.099 | 0.99 |

**Table S11: Post hoc contrasts for the effect of finger condition on the EMG to normalized force ratios for FDS, FDP, and EDC.** Post hoc multiple comparison of means was performed using Tukey method and adjusted $p$-values were obtained by applying the Bonferroni-Holm correction. $n = 16$.

| | | Estimate | Std. Error | df | t ratio | p-value |
|---|---|---|---|---|---|---|
| FDS | Free - 1J | 0.248 | 0.145 | 45 | 1.714 | 0.21 |
| | Free - 2J | 0.422 | 0.145 | 45 | 2.917 | 0.01 |
| | 1J - 2J | 0.174 | 0.145 | 45 | 1.203 | 0.46 |
| FDP | Free - 1J | 0.2077 | 0.107 | 45 | 1.939 | 0.14 |
| | Free - 2J | 0.2804 | 0.107 | 45 | 2.618 | 0.03 |
| | 1J - 2J | 0.0727 | 0.107 | 45 | 0.679 | 0.78 |
| EDC | Free - 1J | 0.588 | 0.169 | 45 | 3.485 | 3e-3 |
| | Free - 2J | 0.692 | 0.169 | 45 | 4.104 | 5e-4 |
| | 1J - 2J | 0.104 | 0.169 | 45 | 0.619 | 0.81 |

**Table S12: Linear regression for testing the effect of $\Delta cc_{EDC}$ on $\Delta f_{max}$ for 16 subjects**. Linear regression with $\Delta cc_{EDC}$ as the explanatory variable and $\Delta f_{max}$ as the dependent variable shows that the effect of $\Delta cc_{EDC}$ on $\Delta f_{max}$ is significant.

Model: $\Delta f_{max} = \alpha + \beta \, \Delta cc_{EDC}$

| | Estimate | Std. Error | t-value | p-value |
|---|---|---|---|---|
| $\alpha$ | 0.159 | 0.049 | 3.258 | 3e-3 |
| $\beta$ | -0.652 | 0.158 | -4.115 | 3e-4 |
| $R^2$=0.36 | | | | |

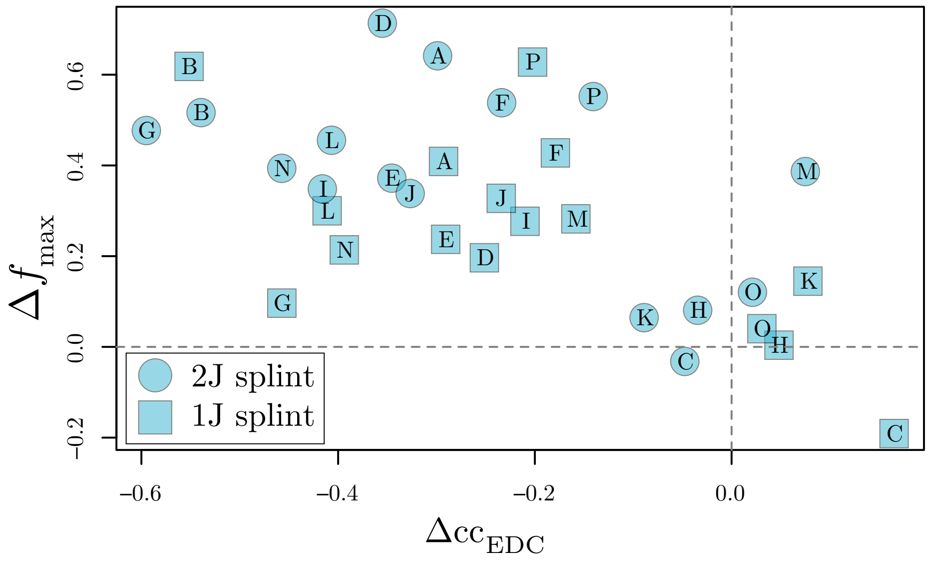

**Fig. S5: Change in $\Delta f_{max}$ as a function of $\Delta cc_{EDC}$** shows that a decrease in EDC contraction is correlated with an increase in the fingertip force upon adding the splint (n=16). See table S12.

**Table S13: Linear regression with $\Delta cc_{EDC}$ as the dependent variable and $F_{free}$ (N) as the explanatory variable** for 16 subjects.

Model: $\Delta cc_{EDC} = \alpha + \beta F_{free}$

| | Estimate | Std. Error | t-value | p-value |
|---|---|---|---|---|
| $\alpha$ | -0.355 | 0.075 | -4.748 | 5e-05 |
| $\beta$ ($N^{-1}$) | 0.004 | 0.002 | 1.819 | 0.08 |
| $R^2$=0.09 | | | | |

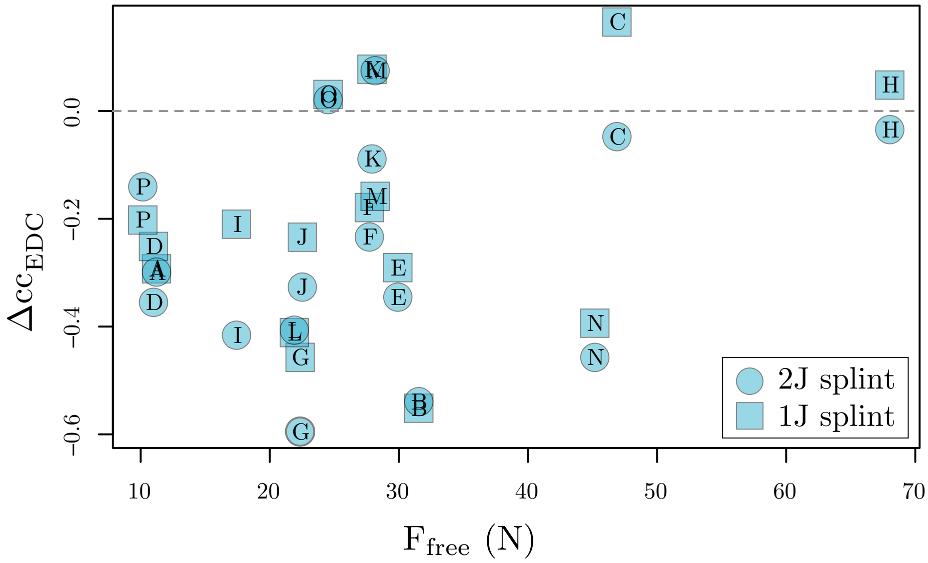

**Fig. S6: Change in $\Delta cc_{EDC}$ as a function of $F_{free}$ (N)** shows that $\Delta cc_{EDC}$ and $F_{free}$ (N) are not correlated (n=16). See table S13.

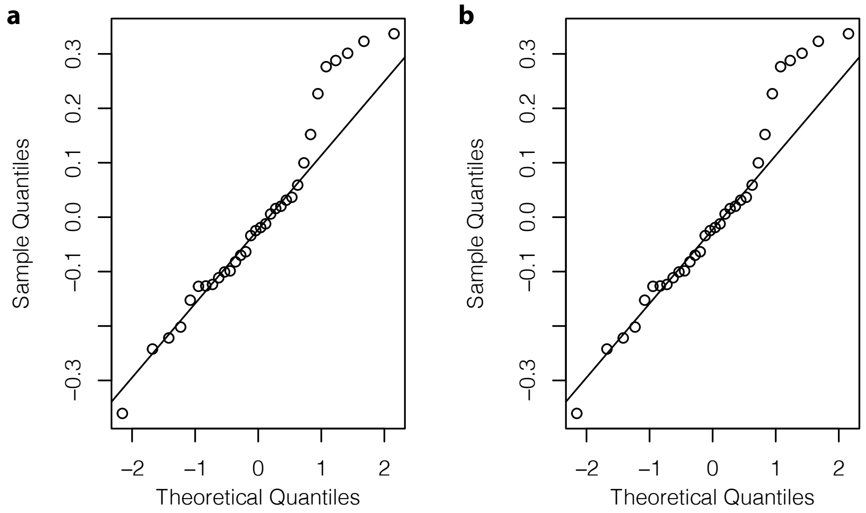

**Fig. S7: Normality of the residuals from the linear models used for regressions related to the co-contraction analysis** in tables (**a,**) S12 and (**b,**) S13.

#### References

[1] Valero-Cuevas, F. J., Zajac, F. E. & Burgar, C. G. Large index-fingertip forces are produced by subject-independent patterns of muscle excitation. *Journal of Biomechanics* **31**, 693–703 (1998).

[2] Venkadesan, M. & Valero-Cuevas, F. J. Effects of neuromuscular lags on controlling contact transitions. *Philosophical Transactions of the Royal Society of London A: Mathematical, Physical and Engineering Sciences* **367**, 1163–1179 (2009).

[3] Strang, G. *Introduction to linear algebra* (Wellesley-Cambridge Press Wellesley, MA, 2016), 5 edn.

[4] Cui, L., Perreault, E. J., Maas, H. & Sandercock, T. Modeling short-range stiffness of feline lower hindlimb muscles. *Journal of Biomechanics* **41**, 1945–1952 (2008).

[5] Grant, M. & Boyd, S. CVX: Matlab software for disciplined convex programming, version 2.1 (2014).

[6] Grant, M. & Boyd, S. Graph implementations for nonsmooth convex programs. In Blondel, V., Boyd, S. & Kimura, H. (eds.) *Recent Advances in Learning and Control*, Lecture Notes in Control and Information Sciences, 95–110 (Springer-Verlag Limited, 2008).

[7] Hajian, A. Z. & Howe, R. D. Identification of the mechanical impedance at the human finger tip. *Journal of Biomechanical Engineering* **119**, 109–114 (1997).

[8] An, K.-N., Chao, E., Cooney, W. & Linscheid, R. Forces in the normal and abnormal hand. *Journal of Orthopaedic Research* **3**, 202–211 (1985).

[9] Holzbaur, K. R., Murray, W. M. & Delp, S. L. A model of the upper extremity for simulating musculoskeletal surgery and analyzing neuromuscular control. *Annals of biomedical engineering* **33**, 829–840 (2005).

[10] Linscheid, R. L., An, K.-N. & Gross, R. M. Quantitative analysis of the intrinsic muscles of the hand. *Clinical Anatomy: The Official Journal of the American Association of Clinical Anatomists and the British Association of Clinical Anatomists* **4**, 265–284 (1991).

[11] Infantolino, B. W. & Challis, J. H. Architectural properties of the first dorsal interosseous muscle. *Journal of Anatomy* **216**, 463–469 (2010).

[12] Jacobson, M. D. *et al.* Architectural design of the human intrinsic hand muscles. *The Journal of hand surgery* **17**, 804–809 (1992).
